# The phenotypic landscape of the mycobacterial cell

**DOI:** 10.1101/2025.11.14.688347

**Authors:** Nadia Herrera, Horia Todor, Lili M. Kim, Hannah N. Burkhart, Evan Billings, Allison Fay, Theodore C. Warner, So Young Lee, Natalie Y. Sayegh, Barbara Bosch, James Chen, Laura L. Kiessling, Michael S. Glickman, Filippo Mancia, Jeremy M. Rock, Carol A. Gross

## Abstract

The Mycobacteriales are an order of diverse bacteria that thrive in many environmental and host-associated niches. Because the most notorious member of this clade, *Mycobacterium tuberculosis,* is a major human pathogen, research on Mycobacteriales has focused on pathogenesis, and, as a consequence, many fundamental aspects of Mycobacterial biology remain understudied. Here, we address this gap by performing a genome-wide CRISPRi chemical genomics screen using a diverse set of >35 antibiotics, detergents, and other anti-microbials predominantly targeting the cell envelope of *Mycobacterium smegmatis,* a saprophytic model Mycobacterium. We highlight new information derived from this screen, including the identification of novel functions for previously uncharacterized conserved and essential genes (in mycolic acid and arabinogalactan synthesis), the discovery of a new drug scaffold/protein target pair, and insights into the mechanism of action of two commonly used antibiotics. These data are also a valuable resource for the mycobacterial research community, as they provide thousands of novel phenotypes for uncharacterized genes and meaningful phenotypic correlations between annotated and uncharacterized genes.

## INTRODUCTION

The Mycobacteriales are a large and diverse order of Actinobacteria including both human pathogens and commensals: many Corynebacterium species are part of a healthy microbiome^1^, and many *Mycobacterium* species are pathogenic. *M. tuberculosis,* the etiological agent of tuberculosis, a disease that kills 1.2 million people annually^2^ and for which ∼¼ of the world’s population is sero-positive^3^. Because of the extreme impact of *M. tuberculosis* as well as *M. leprae* and other non-tuberculosis mycobacterial (NTM) pathogens on human health, much of the research on *Mycobacteria* is understandably disease-focused. Numerous studies have elucidated how *M. tuberculosis* infects humans, persists in the lungs, and evades antibiotics.

However, as a result of this focus, our knowledge about many fundamental cell processes is lacking. For example, many of the more than 600 essential genes in *M. tuberculosis* remain partially or completely uncharacterized^4^. Of these essential *Mycobacterial* genes, ∼400 encode conserved essential genes shared with proteobacteria (e.g., *Escherichia coli*), firmicutes (e.g., *Bacillus subtilis*), and actinobacteria (e.g., *Bifidobacterium breve*). The remainder likely function in uniquely mycobacterial essential processes, such as the synthesis of its unusual and impermeable cell envelope, which is critical for its intrinsic antibiotic resistance and lifestyle.

This envelope consists of highly branched arabinogalactan (AG) and lipoarabinomannan (LAM) whose terminal ends are esterified with mycolic acids that form the inner leaflet of a mycomembrane rich in complex species-specific lipids such as glycopeptidolipids (GPL), trehalose di-mycolates (TDM), trehalose polyphelates (TPP), and phthiocerol dimycocerosates (PDIM)^5^. Despite considerable progress, our knowledge of how these layers are constructed, regulated, and functionalized with proteins^6^ remains at best incomplete.

In its simplest form, chemical genomics involves subjecting genome-wide loss-of-function mutant libraries to a wide variety of stressors and identifying chemical-gene interactions such as drug sensitivity or resistance^7^. Although specific mutant phenotypes can individually be informative, much of the utility of chemical genomics for deciphering gene function arises from considering correlated gene phenotypes across many conditions, and this has been used to decipher the function of many uncharacterized essential and non-essential genes^8–10^. Although mutant libraries of *M. tuberculosis* and other bacteria have been screened across multiple conditions^11^, and compendia of genome-wide fitness experiments exist^12^, three factors hamper the utility of these data for determining gene function. First, these experiments were performed by multiple labs using different lab-adapted strains, experimental techniques, and setups. This introduces batch effects that must be accounted for, thus complicating the interpretation of the results^12^. Second, almost all of these screens were performed using the TnSeq methodology, which cannot query essential genes. Although a CRISPRi screen targeting essential and non-essential genes has been performed^11^, only nine antibiotics were tested, limiting the interpretability of gene-gene phenotypic correlations. Finally, almost all of these phenotypic screens were performed using clinically relevant antibiotics or conditions (e.g., mouse infection), which limits the range of discoverable phenotypes and correlations.

Here, we perform a genome-wide CRISPRi-based chemical genomics screen in the saprophytic mycobacterial model system *Mycobacterium smegmatis* using a wide selection of ∼35 unique chemical stresses focused on the cell envelope. Our screen included the first-line anti-tuberculosis drugs isoniazid and ethambutol, the second-line drug D-cycloserine, and numerous other antibiotics and chemical stressors, including detergents and antiseptics. Using this broad selection of stressors, our screens uncovered a wealth of novel phenotypes and gene-gene phenotypic correlations that yielded powerful insights into Mycobacterial biology. (1) We discovered and characterized MSMEG3144, an essential protein that physically associates with and is required for the function of the branching arabinosyltransferase AftC, (2) uncovered roles for two essential genes in mycolic acid synthesis/export, (3) revealed the role UPF0182-family proteins in membrane protein quality control across Actinobacteria, (4) identified a new drug scaffold/target pair, and (5) refined the mechanism of action of commonly used antibiotics such as ethambutol and D-cycloserine. Our results are an important resource for the mycobacterial research community, demonstrate the utility of our broad chemical-genomics approach, and make significant contributions to our understanding of the mycobacterial cell.

## RESULTS AND DISCUSSION

### *M. smegmatis* CRISPRi chemical-genomics screen yields a robust, high-quality dataset

We performed competitive growth experiments in the presence of sub-MIC concentrations of various stressors using a previously described *M. smegmatis* CRISPRi library encompassing ∼170,000 sgRNAs targeting >99% of genes with multiple knockdown levels^4^. We grew this library to exponential phase, sampled the population for sequencing, induced the CRISPRi system with anhydrous tetracycline (ATc), and added each stressor (Figure 1). Our 40 unique stressors consisted of antibiotics targeting diverse processes (e.g., isoniazid, clarithromycin, ethambutol), as well as chemicals likely to affect or be excluded by the mycomembrane (e.g., detergents, acriflavine, polymyxin B). After 10 doublings of exponential growth (maintained through back dilution), we again sampled the population for sequencing (Figure 1, Methods). We calculated the relative fitness^13^ (RF) of each strain in each condition by comparing its relative abundance (quantified by next-generation sequencing of the sgRNA spacers) at the start and end of each experiment (Figure 1, Methods). RF was reproducible across experimental conditions (median *r* = 0.87, Figure S1A-B) and consistent with previous measurements of the library^4^ (*r* = 0.82, Figure S1C). sgRNAs targeting each gene were separated into low-(0 to 0.33), medium-(0.33 to 0.66), and high-predicted knockdown (0.66 to 1) bins based on their predicted strength^4^. RFs were averaged within bins and used to compute chemical-gene interaction scores (S-scores^14^, Table S1, Table S2, Methods), facilitating the identification of strains with condition-specific growth differences and accounting for the variability in RF measurements. Our final dataset consisted of S-scores for each gene at low-, medium-, and high-knockdown levels across the 71 conditions that passed our quality-control criteria: a ∼12,000 by 71 matrix (Figure 1, Table S2, Methods).

**Figure 1.**
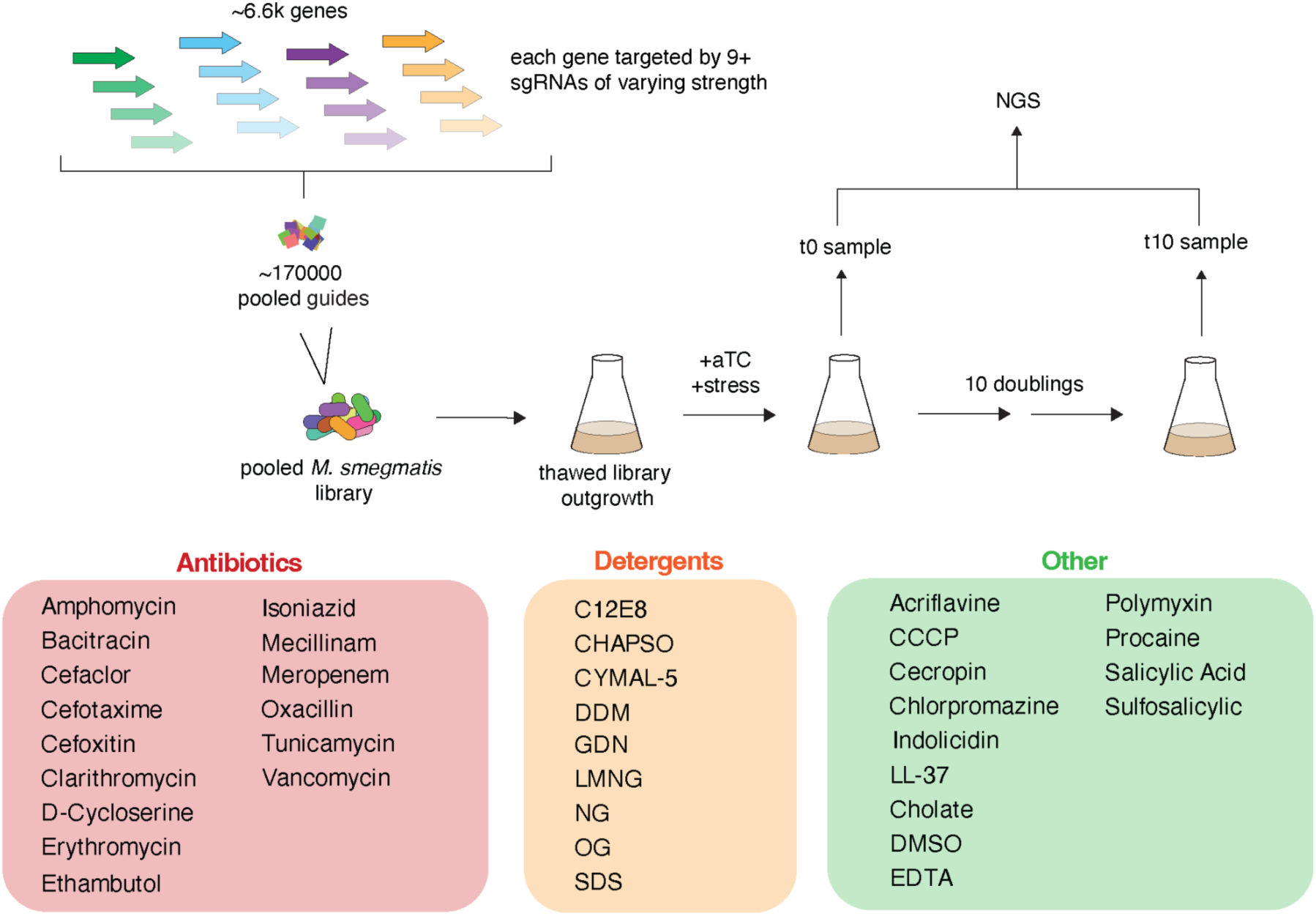
Schematic of the experimental setup. A previously constructed *M. smegmatis* inducible CRISPRi library with ∼170,000 sgRNAs targeting all genes with different levels of knockdown^4^ was used for our chemical-genomic studies. The library was thawed and grown to exponential phase in the absence of inducer. The library was then diluted to an OD600 of 0.05, the CRISPRi system was induced using 100 ng/μL ATc, and the stressor was added. Stressors included diverse antibiotics, detergents, and other chemicals. sgRNA abundance at the start and end of the experiment was enumerated with next-generation sequencing and used to calculate relative fitness (Methods).

*In toto*, our dataset contained 8,169 significant (|S-score| > 5) and 2,148 strong (|S-score| > 7) chemical-gene interactions, as well as many weaker interactions (Figure S1D, Table S2). Gene-chemical interactions identified in our screen reflected many previously reported sensitivities in *M. smegmatis* or closely related species. For example, we confirmed that knockdown of the multidrug efflux pump *lfrA* sensitized cells to acriflavine^15^, that knockdown of *wecA* sensitized cells to tunicamycin^16^, and that knockdown of *inhA* sensitized cells to isoniazid^17^ (Table S2). Our gene-chemical interactions also expanded observations made in other bacteria to *M. smegmatis.* For example, knockdown of *dacB* conferred resistance to vancomycin in *M. smegmatis*, as previously reported for *E. coli* and *B. subtilis*^18^, and knockdown of the putative oleandomycin glycosyltransferase *MSMEG0730* sensitized cells to macrolides (clarithromycin and erythromycin), consistent with its function in *Streptomyces antibioticus*^19^ (Table S2). Importantly, many gene-chemical interactions from our screen reflect new information about gene function or antibiotic mechanisms of action: we found at least one significant phenotype for 45% of genes (Table S2).

### Network of gene-gene interactions reveals connections between core processes and highlights functional partners of genes of unknown function

The chemical interactions (S-scores^14^) of a knockdown strain across all conditions are known as its phenotypic signature, and significant correlations between phenotypic signatures is often indicative of shared function^8^. Consistent with this idea, many of the most highly correlated (r > 0.7, n= 4,009) phenotypic profiles were those between different knockdown levels of the same genes (10.4% of highly correlated interactions), between different genes in the same operon^20^ (due to the polarity of CRISPRi knockdown^21^, 14.0% of highly correlated interactions), or between separately transcribed genes with some experimental evidence of interaction in the STRING database^22^ (9.0% of highly correlated interactions, Figure S2, Table S3).

Phenotypically correlated gene pairs also included many recently discovered interactions that are not in the STRING database. For example, our data showed a high correlation between *ftsZ* and its recently identified transcriptional activator, *ftsR* (*MSMEG3646*)^23^, and between *perM* and the cell division genes *ftsQ* and *ftsL* that co-stabilize its target FtsB^24^ (Table S3). The preponderance of experimentally validated and otherwise expected interactions among correlated gene pairs strongly suggests that high correlations between phenotypic profiles in our data are indicative of shared function and highlights the ability of this approach to uncover new interactions.

To visualize the links between different genes and processes, we constructed a network of gene-gene interactions based on correlated phenotypes (r > 0.7). As expected based on previous work^8,9^, genes with functional relationships clustered together (Figure 2). For example, genes involved in peptidoglycan precursor synthesis (*murABCDEG, ddl*) clustered together near genes involved in peptidoglycan polymerization and modification (*ponA1, pbpA, ftsW, asnB*; Figure 2, inset A). Similarly, genes involved in cytochrome synthesis and maturation clustered together adjacent to *tatC*, which is required for membrane translocation of the Rieske protein encoded by *qcrA*^25^ (Figure 2, inset B). Genes involved in the synthesis of GPL, TPP, or mycolic acids clustered also together, but these mycomembrane constituents as a whole did not, indicating different phenotypes for the depletion of mycolic acids, TPP, GPL, LAM, etc (Figure 2, insets A, C, D). This is strikingly different from what is observed in gram-negative bacteria, where depletion of any outer membrane (OM) associated process (e.g., β-barrel assembly, lipoprotein trafficking, lipopolysaccharide synthesis/export, etc.) compromises the barrier function of the OM and results in similar phenotypes. One interpretation of this finding is that the mycomembrane has a modular composition in which the presence and/or preponderance of various constituent lipids alter properties of the mycomembrane, perhaps in response to environmental conditions or signals, rather than functioning as a unitary permeability barrier.

**Figure 2.**
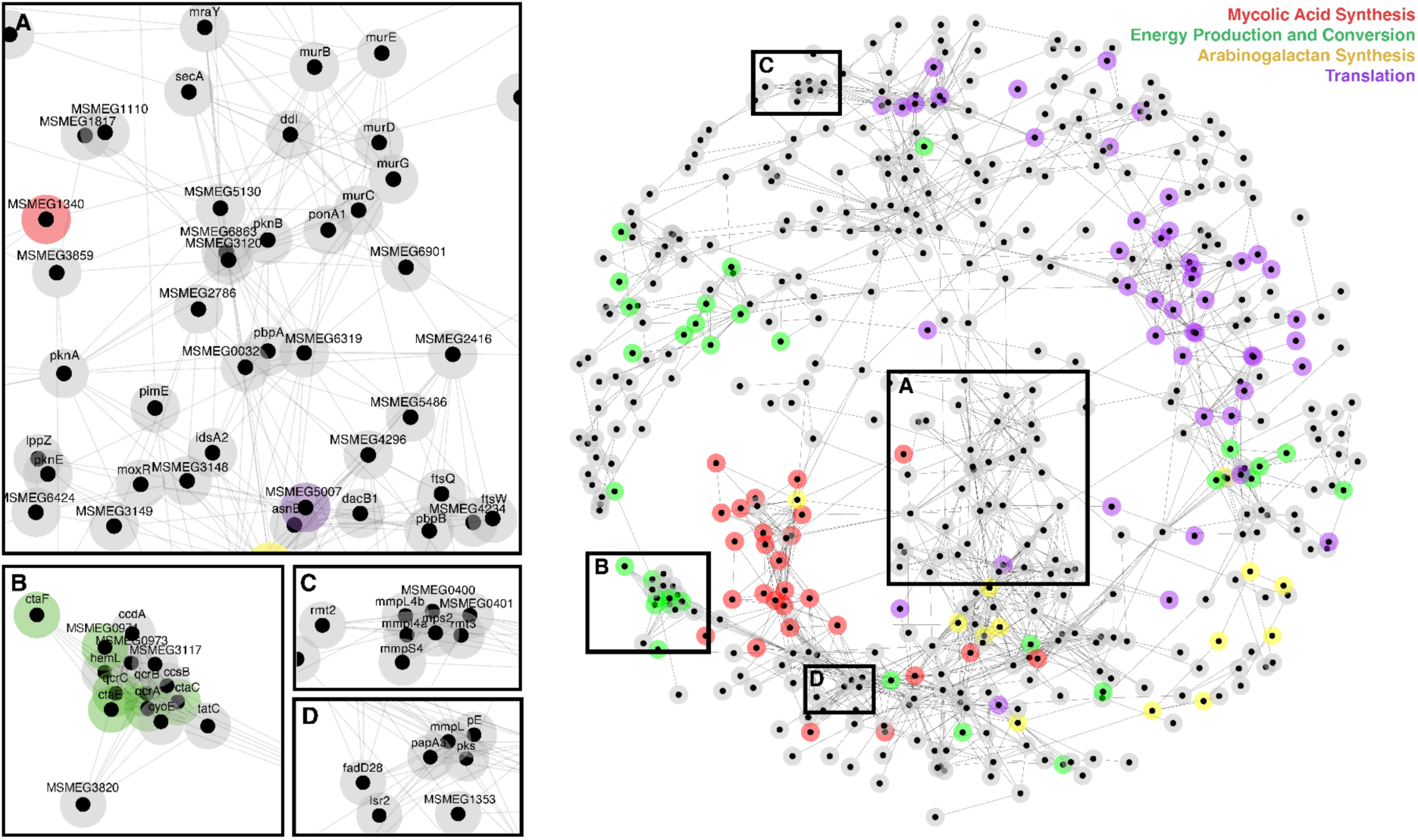
Network of phenotypic correlations. The network was arranged using the edge-weighted spring embedded layout in Cytoscape (Methods), with weight set by correlation strength. Genes outside the main network were excluded. The network includes 480 genes with multiple strong correlations (r > 0.7, total connections: 1523). Genes encoding proteins involved in specific processes clustered together: mycolic acid synthesis (red), energy production and conversion (green), arabinogalactan synthesis (yellow), and translation (purple). Insets highlight processes which cluster together: A) Cluster including genes involved in peptidoglycan precursor synthesis, peptidoglycan polymerization and modification, and LAM synthesis; B) Cluster of genes involved in cytochrome synthesis; C) Cluster of genes involved in GPL synthesis; D) Cluster of genes involved in TPP synthesis.

### Correlated phenotypic profiles identify new players in mycolic acid synthesis/transport

In our network, genes involved in the synthesis and processing of mycolic acids, essential lipids that make up the inner leaflet of the mycomembrane, formed a densely interconnected sub-network (Figure 2, Table S3). The network included genes involved in almost all stages of mycolic acid metabolism, including: synthesis (e.g., *fas, kasA, fadD32-pks13-accD4, accD5*), transport (*ttfA, mmpL3*), and polar positioning (*MSMEG0311*, *MSMEG0315*, *MSMEG0317*, *MSMEG0319*). Ag85 mycolic acid transferases, which are non-essential^4^ due to partial redundancy, were absent. Two genes not previously implicated in mycolic acid synthesis (and whose phenotypes cannot be explained by polar effects) clustered with mycolic acid synthesis: *MSMEG6143* and *MSMEG0834* (Table S3).

To determine whether knockdown of *MSMEG0834* or *MSMEG6143* affects mycolic acid metabolism, we labeled these knockdown strains with AzLac, a recently described mycolic acid mimic that reports on both the location and amount of endogenous mycolic acid conjugation to the arabinogalactan^26^. After labeling with a fluorophore, we quantified AzLac accumulation/cell using flow-cytometry (Methods). As expected, knockdown of the sole mycolic acid transporter *mmpL3* eliminated AzLac labeling, whereas wildtype cells exhibited significant labeling (Figure 3A). Labeling was reduced in both the *MSMEG0834* and *MSMEG6143* knockdown strains compared to the wild-type control, suggesting that these genes directly or indirectly affect mycolic acid synthesis/export. Knockdowns of *MSMEG0834* exhibited less labeling than *MSMEG6143*, but more than *mmpL3*, an order consistent with the relative essentiality of those genes in both *M. smegmatis* and *M. tuberculosis*^4^.

**Figure 3.**
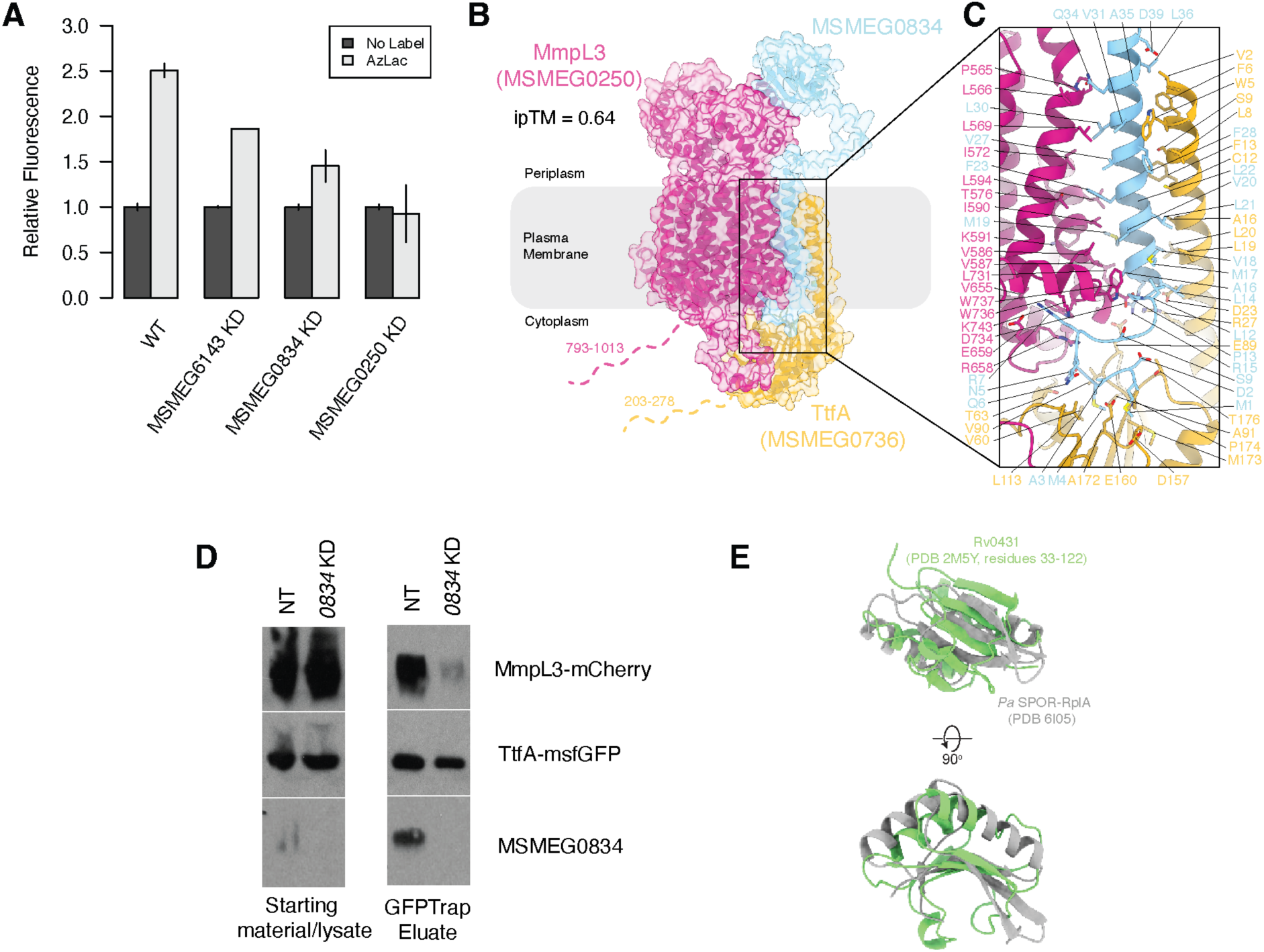
MSMEG0834 and MSMEG6143 are involved in mycolic acid synthesis and/or transport. A) AzLac labeling data show decreased labeling of AG-bound mycolic acid in the *MSMEG6143*, *MSMEG0834,* and *mmpL3* (*MSMEG0250*) knockdown strains compared to WT. Error bars represent the standard deviation of 2 biological replicates. B) AlphaFold3 prediction of the MmpL3 (violet), TtfA (gold), and MSMEG0834 (cyan) complex rendered as cartoons superimposed on a transparent molecular surface. For clarity, the AlphaFold3 predicted C-termini of MmpL3 (residues 793-1013) and TtfA (residues 209-276) were omitted and instead indicted with dashed-lines. C) Zoom-in of boxed area in (B) showing predicted interacting residues TM helices of MmpL3, TtfA, and MSMEG0834, depicted as sticks. Colors are the same as in (B). D) MSMEG0834 is required for the interaction between TtfA and MmpL3. Yield of MmpL3 from a TtfA-msfGFP pulldown is decreased in a MSMEG0834 CRISPRi knockdown strain compared to a nontargeting CRISPRi strain. E) LytR_C domains show structural similarity to SPOR domains. Overlay of the experimental structure of the LytR_C domain of Rv0431 (light green, PDB 2M5Y) and the experimental structure of the SPOR domain of *Pseudomonas aeruginosa* (*Pa*) RplA (grey, PDB 6I05). Structural alignment was performed in PyMOL using ‘cealign’ command (5.81 Å over 64 residues).

*MSMEG6143* encodes a membrane protein with nine transmembrane (TM) helices and a C-terminal periplasmic domain. The protein shares some structural similarity with glycosyltransferases of the GT-C subfamily^27^ and was identified by distant homology searches as an N-oligosaccharyl transferase^28^. We did not pursue further characterization of this gene.

*MSMEG0834* encodes a widely conserved tuberculin-related peptide with a LytR_C domain (Interpro: IPR027381, PFAM: PF13399) and a single TM helix. Its *M. tuberculosis* homolog, Rv0431, has been reported to play a role in the formation of membrane vesicles and immune modulation in *M. tuberculosis*^29,30^, roles consistent with a function in mycolic acid synthesis or transport^31^. MSMEG0834 is essential (Figure S3A), and has been identified in pulldowns of TtfA (trehalose mono-mycolate transport factor A, MSMEG0736/Rv0383c) and MmpL3 (the mycolic acid exporter) in *M. smegmatis*^32^. To better understand how MSMEG0834 may interact with MmpL3 and TtfA, we used AlphaFold3^33^ to co-fold MmpL3, TtfA, and MSMEG0834 (Figure 3B). In the predicted high-confidence complex (ipTM = 0.64), the N-terminal TM helix of MSMEG0834 appears to mediate the interaction between the TM helix of TtfA and the C-terminal TM helices of MmpL3 (Figure 3C), suggesting that MSMEG0834 may be required for MmpL3 to interact with TtfA, its essential partner^32^.

To assess this prediction, we first tested whether the N-terminal helix of MSMEG0834 was sufficient to support growth in the absence of WT-MSMEG0834. We found that the first 49aa (but not the first 43aa) of MSMEG0834 were sufficient to rescue growth, consistent with the hypothesis that only the N-terminal TM helix is essential (Figure S3B). Next, using a previously described^34^ bacterial two-hybrid assay (BACTH), we showed that MSMEG0834 directly interacts with TtfA, while MmpL3 does not (Figure S3C). This finding supports a role for MSMEG0834 in mediating the interaction between MmpL3 and TtfA. Finally, we tested whether MSMEG0834 depletion disrupted the interaction between MmpL3 and TtfA by using GFP-tagged TtfA to pulldown mCherry-tagged MmpL3. Under normal conditions, mCherry-tagged MmpL3 co-eluted with GFP-tagged TtfA from a GFPTrap column (Figure 3D; Methods).

However, in an *MSMSEG0834* CRISPRi strain grown under inducing conditions for 24hrs, mCherry-tagged MmpL3 no longer co-elutes with GFP-tagged TtfA, suggesting that MSMEG0834 is crucial for the interaction of these two essential proteins (Figure 3D). Together, these data support the idea that the N-terminal TM helix of MSMEG0834 mediates the interaction between MmpL3 and TtfA and suggest that the essentiality of MSMEG0834 may be due to this role.

The apparent dispensability of the LytR_C domain of MSMEG0834 raises questions about its function. LytR_C domains are predominantly found in actinobacteria and frequently associated with LCP proteins^35^, which transfer secondary cell wall polymers to PG. Intriguingly, we found weak but significant structural similarity between LytR_C and SPOR domains despite low sequence similarity and different electrostatic profiles (Foldseek^36^ e-values: 5.8e^-7^ to 1.1e^-5^ between the *M. tuberculosis homolog of* MSMEG0834, Rv0431 (2M5Y^29^), and SPOR domains; Figure 3E; Figure S3D). In proteobacteria and firmicutes, SPOR domains localize cell division proteins by binding denuded septal PG^37^. It is tempting to speculate that LytR_C domains function analogously to SPOR domains (which are rare in actinobacteria), recruiting specific proteins (e.g., LCPs, MmpL3) to the septum by binding actinobacterial-specific septal PG epitopes, though this remains to be demonstrated.

Together, our studies implicate MSMEG0834 and MSMEG6143 in mycolic acid synthesis/export/regulation and suggest that MSMEG0834 functions in a complex with MmpL3 and TtfA.

### MSMEG3144 forms a complex with AftC and is required for AftC activity

*embA*, *embB*, *aftB*, and *aftC* cluster together in our gene-gene network (Figure 2, Table S3), consistent with their shared function in AG synthesis. Interestingly, *MSMEG3144*, encoding a DUF6676 family protein, also clustered with these genes, exhibiting a particularly strong correlation with *aftC* (*r* = 0.90) (Figure 4A-B). *aftC* and *MSMEG3144* homologs (*cgp_2077* and *cgp_1736*) were also highly and specifically correlated (r = 0.89, Figure S4A-B) in TnSeq based chemical genomics experiments performed in the distantly related mycolic acid-producing bacteria *Corynebacterium glutamicum*^38^, suggesting that the phenotypic similarity between depleting/inactivating *aftC* and *MSMEG3144* was not due to our choice of conditions or to our use of CRISPRi, but may reflect an evolutionarily conserved functional relationship.

**Figure 4.**
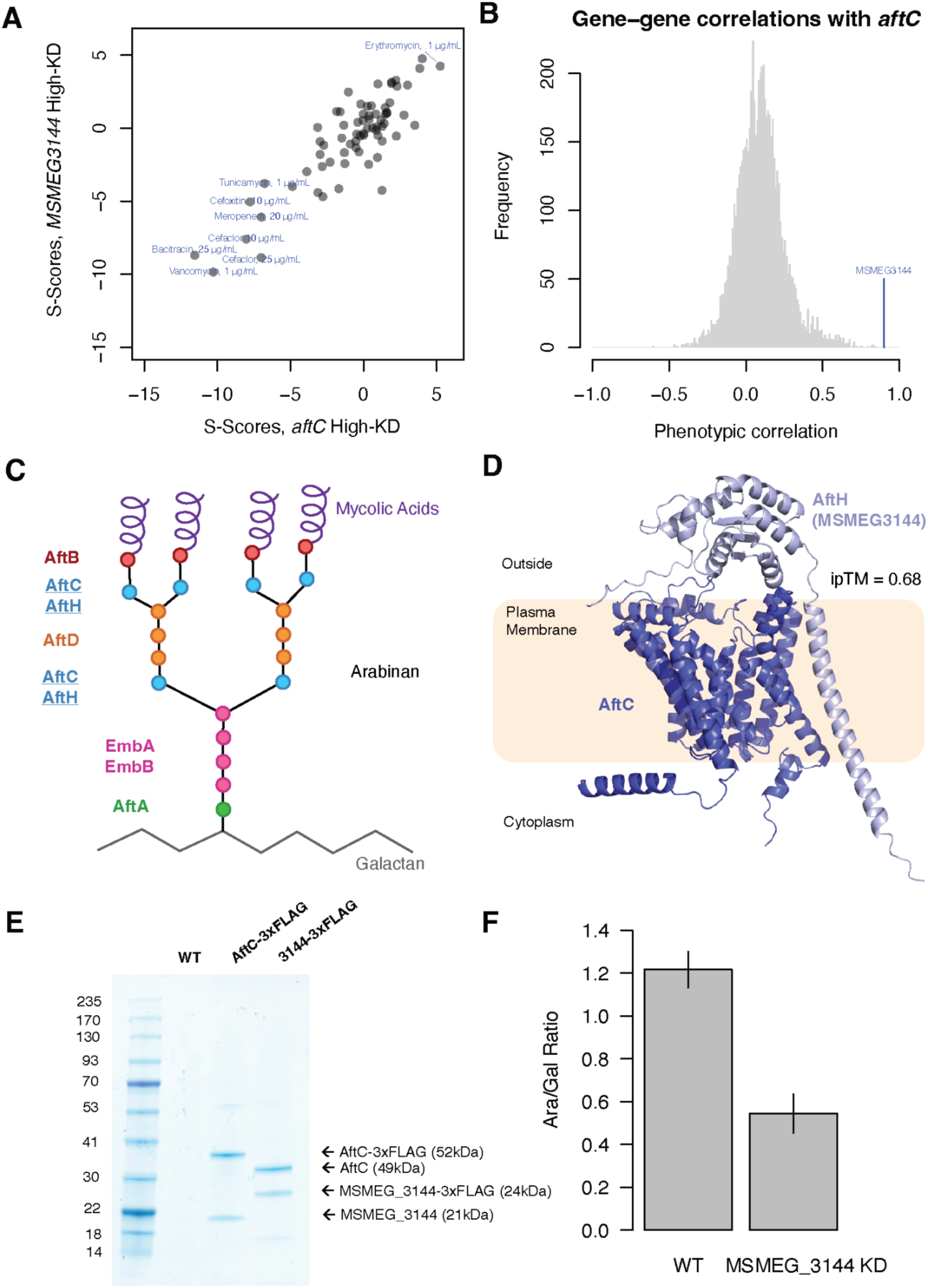
MSMEG3144 is an essential partner of AftC. A) S-scores of *aftC* and *MSMEG3144* (from the high-knockdown bins) in various conditions. Conditions where one or both genes have |S-score| > 5 are labeled. B). Histogram of correlation values of all genes with *aftC*. C) Schematic of arabinofuranosyl transferase activity in Mycobacteria. Each circle represents an arabinofuranose residue, and circle color indicates the enzyme that appends each residue according to ref^82^. AftC and AftH (MSMEG3144) are underlined. D) High-confidence (ipTM = 0.68) AlphaFold3 predicted structure of AftC and AftH (MSMEG3144). E) MSMEG3144 and AftC physically interact. Coomassie stain of WT, AftC-3xFLAG, and MSMEG3144-3xFLAG solubilized membrane fractions after purification. Using ORBIT, a payload plasmid containing a 3xFLAG was inserted in the genomic region flanking *MSMEG2785* (*aftC*) and *MSMEG3144* (Figure S4E). Membranes of the AftC-3xFlag and MSMEG3144-3xFLAG *M. smegmatis* cells were solubilized and purified using α-FLAG resin. The elution fractions of WT, AftC-3xFLAG, MSMEG3144 cells were analyzed with SDS-PAGE. When tagged, both AftC and MSMEG3144 reciprocally co-purify with the other. F) The arabinan/galactan ratio decreases in cells with *MSMEG3144* knockdown. Error bars represent the standard deviation of 2 biological replicates.

Knockdown of *MSMEG3144* caused growth and morphological defects (Movie S1-2), and these morphological defects were distinct from those of knocking down *MSMEG3145* (*ripA*), a well characterized peptidoglycan hydrolase encoded adjacent to MSMEG3144 (Movie S3). The growth defect of the *MSMEG3144* knockdown strain could be rescued by expressing a CRISPRi-resistant complementation construct carrying *MSMEG3144* (Figure S4D). Together, these data confirm the fitness defect of the *MSMEG3144* strain observed in the pooled screen, eliminate the possibility that it is due to a polar effect, and implicate MSMEG3144 in some aspect of cell wall synthesis, consistent with its phenotypic correlation to the *aftC* knockdown strain.

The branched arabinan of AG is assembled by the action of seven essential arabinofuranosyltransferases^39^ (AftA-D, EmbABC), each of which transfers arabinose from the lipid-linked donor decaprenylphosphoryl arabinofuranose (DPA) onto specific acceptor sugars (Figure 4C). To determine if MSMEG3144 may function in a complex with one of these proteins, we co-folded each of the seven arabinofuranosyltransferases with MSMEG3144 and assessed the interface score (ipTM) of the predicted complex (Figure S4C). Only the MSMEG3144-AftC complex exhibited a high ipTM (0.68), whereas the other arabinofuranosyltransferases exhibited low ipTMs (< 0.4). The high-confidence interaction between MSMEG3144 and AftC was also conserved for the *M. tuberculosis* and *C. glutamicum* homologs of these genes (Figure S4C). In the predicted structure of the MSMEG3144-AftC complex, the defined TM helix of MSMEG3144 associated with AftC via interactions with its TM helices, and the MSMEG3144 periplasmic domain made additional interactions with the periplasmic loops of AftC, suggesting that MSMEG3144 may play a structural role similar to the CTD of the other arabinofuranosyltransferases^40–43^, capping the active site of AftC and positioning the correct substrate for catalysis (Figure 4D). The structure of the MSMEG-3144-AftC complex makes two predictions: 1) that MSMEG3144 and AftC physically associate, and 2) that both proteins are needed for AftC function.

To test whether AftC and MSMEG3144 interact *in vivo*, we used ORBIT^44^ to construct strains with AftC and MSMEG3144 3xFLAG tagged at their endogenous loci (Figure S4E) and performed reciprocal pull-down assays. We found that MSMEG3144 and AftC specifically co-purified with each other, indicating a strong *in-vivo* interaction (Figure 4E).

To test whether MSMEG3144 is required for the enzymatic function of AftC, we analyzed the composition of the *M. smegmatis* cell wall in the *MSMEG3144* knockdown strain (Methods).

AftC catalyzes the addition of α(1→3) arabinofuranose residues to AG, introducing branch points in the arabinan polymer^45^. Therefore, depleting AftC results in a drastically decreased arabinose/galactan (Ara/Gal) ratio^45^. We reasoned that, if MSMEG3144 is required for AftC function, its depletion should also affect the Ara/Gal ratio. Consistent with this hypothesis, the *MSMEG3144* knockdown strain exhibited an Ara/Gal ratio of ∼0.5, which is consistent with what has been reported for an *aftC* deletion (0.4)^45^ and significantly lower than what was reported for *embA, embB*, or *embC* insertional inactivation mutants (0.88-1.41)^46^ (Figure 4F). By contrast, the wildtype strain exhibited an Ara/Gal ratio of ∼1.2, which is lower than but broadly consistent with previous studies^45,46^. These results suggest that MSMEG3144 is required for AftC function and may explain the low observed activity of purified AftC^47^. Taken together, our phenotypic, biochemical, and structural analyses suggest that MSMEG3144 functions as an essential partner of AftC in evolutionarily diverse mycolic acid-producing bacteria, and we propose that MSMEG3144 and its homologs be renamed AftH (for <u>a</u>rabinofuranosyltransferase <u>h</u>elper).

### UPF0182-family proteins are involved in the regulation of FtsH-mediated proteolysis in Actinobacteria

UPF0182-family proteins are universally conserved in Actinobacteria and have been associated with diverse phenotypes (e.g., adhesion in *Corynebacterium diphtheriae*^48^ or iron homeostasis in *M. tuberculosis*^49^). They generally consist of 7 TM helices and a large periplasmic domain. In our screen, knockdown of *MSMEG1959* (the sole UPF0182-family protein in *M. smegmatis*) resulted in sensitivity to Polymyxin B and resistance to erythromycin and clarithromycin (as well as other phenotypes, Table S2). These distinct phenotypes were strongly correlated to those of *MSMEG4237* (a small conserved transmembrane protein, r = 0.87), *ftsH* (a ubiquitous membrane-bound ATP-dependent protease, r = 0.80), and *MSMEG5275* (*yccA* - a regulator of FtsH activity, r = 0.67, Table S3). The phenotypic link between UPF0182-family proteins and FtsH/YccA is not restricted to *M. smegmatis.* TnSeq chemical genomics experiments in *C. glutamicum* also show a strong correlation between the phenotypes of the UPF0182-family homolog (*cgp_0896)* and *ftsH* (r = 0.80) or the *yccA* homolog (r = 0.77)^38^. Moreover, even in the distantly related actinobacterium *Bifidobacterium breve*, TnSeq experiments show a substantial similarity between the phenotypes of the UPF0182-family homolog and the YccA homolog (r = 0.62, reciprocal top hits^50^) (Figure 5A).

**Figure 5.**
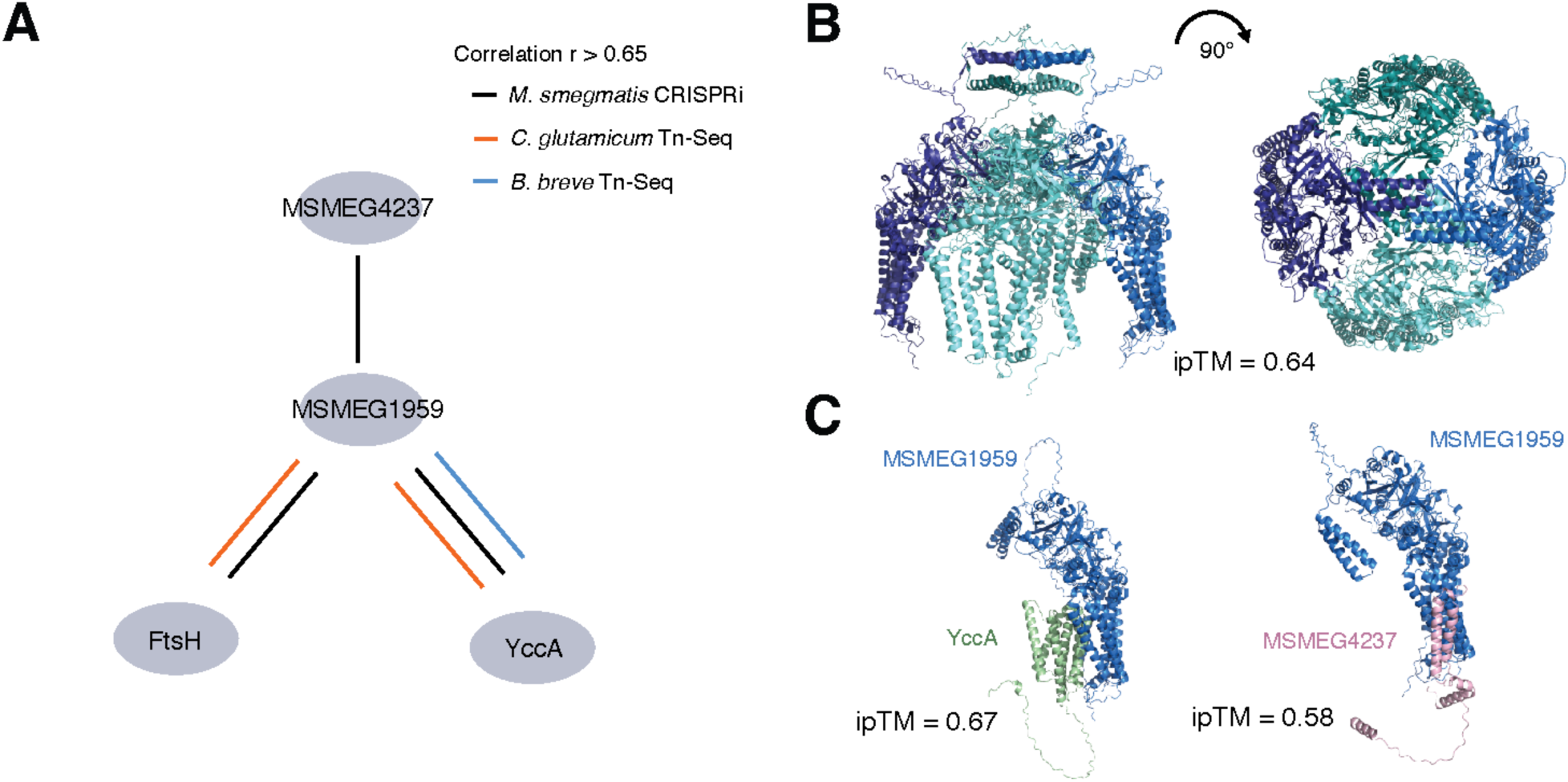
A) UPF0182-family proteins (exemplified by *M. smegmatis* MSMEG1959) are phenotypically correlated to FtsH and/or its regulator YccA in *M. smegmatis* as well as in the evolutionarily distant actinobacteria *C. glutamicum,* and *B. breve.* B) AlphaFold3 predicted structure of the MSMEG1959 homo-tetramer, showing the periplasmic dome supported by TM helices. C) Predicted binding of YccA and MSMEG4237 to MSMEG1959.

In *E. coli,* FtsH is responsible for general membrane protein quality control but is also involved in the regulation of multiple essential processes by degrading key proteins: LpxC in lipopolysaccharide synthesis^51^, SecY in protein secretion^52^, and RpoH in the heat-shock response^53^. YccA, a homolog of the eukaryotic Bax Inhibitor-1, can protect SecY from FtsH degradation when SecY is jammed by mis-targeted proteins or translation-inhibiting antibiotics^52^. The phenotypic association of UPF0182 family proteins with FtsH and YccA strongly suggests a role for this protein family in the regulation of FtsH.

To better understand how MSMEG1959 may work with FtsH and/or YccA, we predicted the structure of MSMEG1959 using AlphaFold3^33^. MSMEG1959 appears to be a tetramer (based on the higher ipTM score of the tetramer (0.64) compared to the dimer (0.22) or trimer (0.49). Tetrameric MSMEG1959 is predicted to form a dome, with contacts between the periplasmic domains (Figure 5B). The superficial similarity of this structure to the nautilus-like HflC/K complex that regulates FtsH in *E. coli*^54^ initially led us to ask whether FtsH could bind inside, but structural modeling did not support this possibility. We next tested whether MSMEG1959 was predicted to bind YccA and/or MSMEG4237. MSMEG1959 was predicted to interact with both YccA (inside the dome, ipTM = 0.67, Figure 5C) and MSMEG4237 (outside the dome, ipTM = 0.58, Figure 5C). Together, these data suggest that the shared phenotypes of *MSMEG1959*, *MSMEG4237*, and *yccA* knockdown strains are due to a physical interaction and implicate these proteins in FtsH-associated processes in Actinobacteria. A role for MSMEG1959 in membrane protein degradation is also consistent with its extensive conservation and with the diversity of phenotypes reported for deletion/depletion mutants of this gene in different actinobacteria.

### Refining the mechanism of common antimicrobials and revealing potential synergies

Antimicrobials and antibiotics function primarily by targeting essential processes. Since CRISPRi depletion of an antibiotic’s target is often sensitizing^9^, our CRISPRi screen can reveal and refine the mechanism of antibiotic action, as we demonstrate below for two commonly used drugs.

#### D-cycloserine likely affects D-glutamate synthesis through D-alanine-transaminase

D-cycloserine (DCS) is a broad-spectrum antibiotic and part of the core second-line treatment group B listed in the WHO guidelines for treatment of multidrug and extensively drug-resistant-TB (MDR/XDR-TB). Its primary activity is the inhibition of D-ala-D-ala ligase (*ddl*) and the pyridoxal 5-phosphate (PLP) dependent enzyme alanine racemase (*alr*), which together produce the D-ala-D-ala moiety of the peptidoglycan stem peptide^55^. Our screen identified additional sensitizing hits: *murI* (glutamate racemase, which produces D-glutamate for PG synthesis; *MSMEG4903*), *thyX* (folate-dependent thymidylate synthase; *MSMEG2683*), and *MSMEG4904* (likely due to polar knockdown of *murI*^21^) (Figure S5, Table S2). Neither ThyX nor MurI are known to be directly inhibited by DCS, but the sensitivity of *murI* to DCS can be explained by the presence of D-alanine transaminase, a PLP-dependent enzyme that couples D-glu and D-ala synthesis^56,57^ (Figure 6A). Consistent with this hypothesis, transposon insertion mutants of *murI* in diverse bacteria that encode D-alanine transaminase are also strongly and specifically sensitized to DCS (Figure 6B)^10^. It remains unclear if this phenotype is due to inhibition of D-alanine transaminase (as reported for the *Aminobacterium colombiense* enzyme^58^), inhibition of Alr (which would decrease the availability of D-ala), or both.

**Figure 6.**
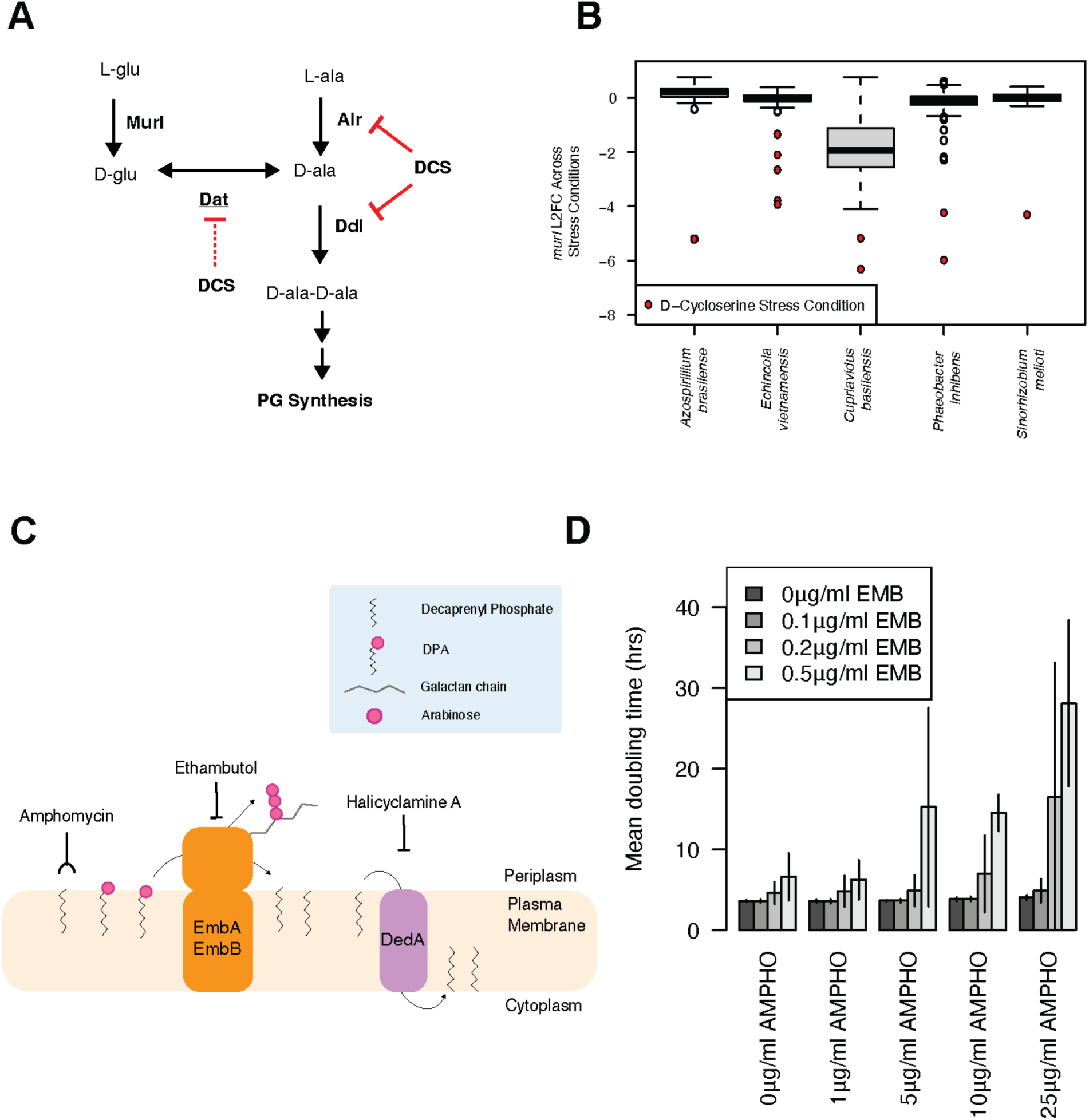
CRISPRi chemical-genomics refines the mechanism of action of two antibiotics. A) Schematic of the D-alanine and D-glutamate synthesis pathway and its inhibition by DCS. Inhibition of Alr and Ddl is depicted using solid lines, since it is established in the literature. Novel inhibition of D-alanine transaminase is shown in dotted lines. B) *murI* mutants of diverse bacteria are sensitive to DCS (from ref^10^). C) Schematic of AG elongation by EmbAB and decaprenylphosphate recycling by DedA. D) Ethambutol and amphomycin synergize. Error bars represent the standard deviation of 2 biological replicates.

Independent of mechanism, these observations explain the previously surprising finding that the glutamate racemase inhibitor β-Chloro-d-Alanine^59^ and DCS exhibit extreme synergy against *M. tuberculosis*^60^. The sensitivity of *thyX*, which encodes the essential thymidylate synthase of mycobacteria, to DCS remains puzzling but may reflect a previously hypothesized second essential function of ThyX^61^.

#### Ethambutol treatment may sequester decaprenyl-phosphate

Ethambutol is a first-line antituberculosis drug that, in *M. smegmatis*, acts by inhibiting the essential arabinosyltransferases EmbA and EmbB^62^. In our screen, knockdown strains targeting *embA* (*MSMEG6388*) and *embB* (*MSMEG6389*) were highly sensitized to ethambutol, as expected. Strikingly, knockdown of *MSMEG0750*, which encodes the sole and essential DedA-family decaprenyl-phosphate (DP) flippase of *M. smegmatis*^63^, also strongly sensitized cells to ethambutol (Figure S6A, Table S2), suggesting that a secondary effect of ethambutol is the sequestration of DP, which is also required for peptidoglycan synthesis, into DPA, analogous to what has been described when interrupting ECA or O-antigen synthesis in *E. coli*^64^. This phenomenon is not limited to *M. smegmatis*: ethambutol sensitivity is also observed for knockdown strains targeting *dedA* (*rv0364*) in *M. tuberculosis*^11^. Decaprenyl phosphate sequestration is expected to be strongest for EmbA/B, since these enzymes typically use many DPA donors to extend the arabinan chain (Figure 6C). To test whether lipid sequestration is a relevant secondary effect of ethambutol, we tested whether ethambutol synergizes with amphomycin, an antibiotic that sequesters decaprenyl-phosphate on the outer leaflet of the inner membrane. Whereas neither 2 µg/ml of ethambutol nor 10-25 µg/ml of amphomycin drastically affects the growth of *M. smegmatis* (doubling time 4-5hrs), in combination the two drugs almost completely blocked growth (doubling time >15hrs, Figure 6D), indicating a strong synergistic effect. Similar, but less dramatic synergies were observed for other concentrations of ethambutol and amphomycin. No synergy was observed between 2 µg/ml of ethambutol and either erythromycin or rifampicin, suggesting that the observed synergy between ethambutol and amphomycin is specific and not due to membrane permeabilization by ethambutol (Figure S6B-C). These results suggest that combining ethambutol with a drug that antagonizes DedA, such as amphomycin^65,66^ or halicyclamine A^67,68^, is likely to be a viable strategy for enhancing ethambutol effectiveness.

Our findings refine the well-established mechanisms of DCS and ethambutol and demonstrate the utility of our CRISPRi method for understanding drug action.

### Salicylic acid and PanB are an evolutionarily conserved scaffold-target combination

Salicylic acid (SA) is a known antimicrobial^69^ without a precise molecular mechanism. *M. smegmatis* produces SA as a precursor to mycobactin siderophores; however, it is toxic at higher concentrations^70^. In our CRISPRi screen, knockdown strains targeting the early steps of coenzyme A (CoA) synthesis (*panB-MSMEG4298*, *panC-MSMEG6097*, *panG-MSMEG6098*, *coaA-MSMEG5252*) were sensitized to SA, raising the possibility that SA antagonizes one of these enzymes (Figure 7A). Consistent with the idea that SA limits CoA synthesis, several TCA cycle enzymes such as *sucCD* (succinate-CoA ligase) and *gltA* (citrate synthase) were also sensitized (Figure S7A, Table S2). To test this hypothesis, we constructed a *panB* CRISPRi strain and tested its susceptibility to SA (Methods). In the absence of SA, the *panB* CRISPRi strain grew as well as wild-type (WT) over 24 hours, likely due to excess CoA and PanB transcribed and/or translated before the onset of CRISPRi repression. However, the *panB* strain was exquisitely sensitive to SA (Figure 7B), exhibiting reduced growth at SA concentrations that did not affect WT. The *panB* strain did not exhibit sensitivity to either propionic acid or acetic acid (Figure S7B-C), suggesting that its sensitivity to SA is not a general sensitivity to weak acids or protonophores. The sensitivity of the *panB* strain to SA was rescued by the addition of 24 µg/ml pantothenate to the media, confirming the action of SA against this pathway. These results suggest that salicylic acid inhibits CoA synthesis upstream of pantothenate.

**Figure 7.**
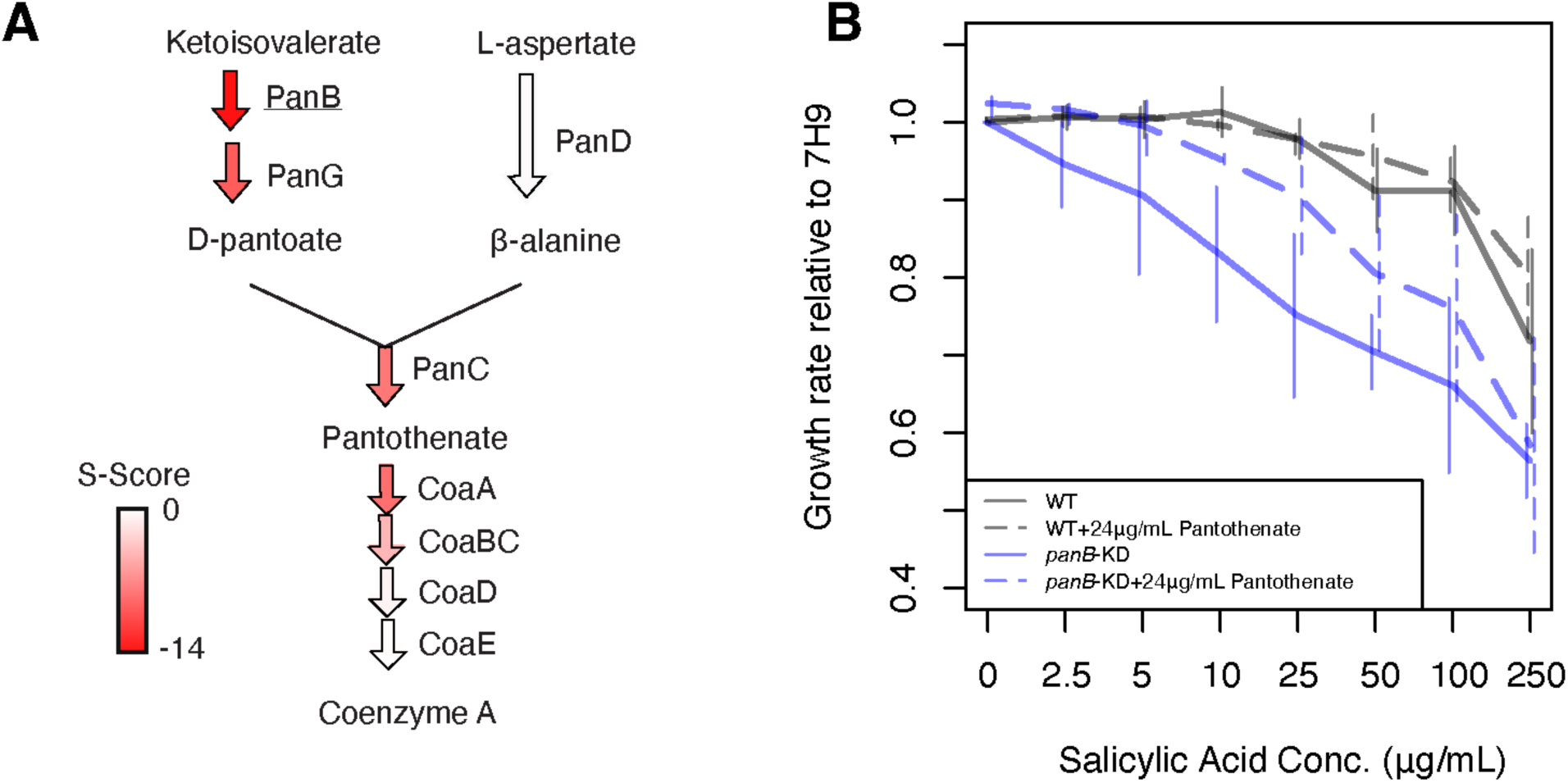
Salicylic acid antagonizes PanB. A) Schematic of the pathway of CoA synthesis, with the S-score of each gene represented by the color of the arrow. PanB is the most sensitized, and sensitivity decreases in the later steps of the pathway. B) The *panB* knockdown strain is sensitized to SA (solid blue lines), and the addition of exogenous pantothenate (dashed blue lines) antagonizes the effect. No change is seen for WT (solid black lines) upon pantothenate supplementation (dashed black lines). Error bars represent the standard deviation of 2-5 biological replicates.

Several *E. coli* studies published before the advent of molecular genetics support this hypothesis^71,72^. These studies identified a concentration range in which the growth-inhibitory effects of SA on WT *E. coli* could be antagonized by the addition of exogenous CoA precursors, including pantoate and pantothenate. They used this data to conclude that salicylic acid antagonizes pantoate synthesis. Pantoate is synthesized from α-ketoisovalerate by the sequential actions of PanB and either PanE or PanG (Figure 7A). PanB is a widely conserved enzyme, with substantial sequence homology even between *E. coli* and *M. smegmatis* (∼46% identity), whereas PanE (the *E. coli* 2-dehydropantoate 2-reductase) and PanG (the *M. smegmatis* 2-dehydropantoate 2-reductase) share no significant homology^73^. Thus, the most parsimonious explanation for SA growth inhibition in both *E. coli* and *M. smegmatis* is that SA antagonizes PanB.

PanB is a TIM-barrel enzyme that transfers a hydroxymethyl group from 5,10-methylenetetrahydrofolate (mTHF) to alpha-ketoisovalerate to form ketopantoate. Drugs with similarity to the para-aminobenzoic acid moiety of folate such as para-aminosalicylic acid or sulfonamides act as folate antimetabolites^74,75^, raising the possibility that SA antagonizes PanB through a folate-related mechanism, including (but not limited to) direct competition with mTHF for binding to PanB. Taken together, our data establish SA-PanB as an evolutionarily conserved scaffold-target pair. A PanB inhibitor based on SA will likely have high selectivity for bacteria (since humans don’t synthesize pantothenate) and it may synergize with the first-line anti-tuberculosis drug pyrazinamide (which has been suggested to target PanD^76^).

### PERSPECTIVE

Mycobacteria have been almost exclusively studied from the perspective of developing clinically relevant drugs and strategies to combat *M. tuberculosis* and other pathogenic Mycobacteria.

This focus has resulted in innovative drugs and therapies against *M. tuberculosis*, but has also contributed to a limited understanding of Mycobacterial, and by extension, Actinobacterial biology. We addressed this shortcoming by performing a broad chemical genomics screen in *M. smegmatis.* Our screen differed from previous screens in three important ways: 1) we used CRISPRi, instead of transposon mutagenesis, allowing us to measure the chemical interactions of both essential and non-essential genes; 2) each gene was targeted by multiple sgRNAs of different predicted efficacy, increasing reproducibility and the chance of observing relevant phenotypes and; 3) we selected a broad spectrum of antibiotics, antiseptics, and chemical stressors to elicit phenotypes from distinct subsets of genes, increasing the interpretability of correlated phenotypic profiles. Our analysis of these results exemplifies the kinds of information that can be gleaned from this approach.

First, our screen uncovered phenotypic correlations between genes of unknown function and well-characterized partners. These connections suggested targeted experiments that allowed us to assign functions to 3 essential genes of unknown function. We identified a phenotypic correlation between *aftC* and the essential gene *MSMEG3144* (*aftH*), suggesting that these genes may work together. AlphaFold3 co-folding of these two proteins indicated that they form a complex, and that this complex is required for the function of AftC: hypotheses that we experimentally validated. Though the structural mechanism underlying these observations remains to be elucidated, essential activators have been identified for some GT-C enzymes involved in GPI-anchor synthesis in eukaryotes^77,78^, suggesting that “bipartite” GT-C like AftC/AftH may be a common theme in all domains of life. Strong phenotypic correlations also helped us link MSMEG0834 and MSMEG6143 to mycolic acid synthesis or transport: a hypothesis we supported by using a recently reported mycolic acid probe to show that knockdown of these genes affects mycolic acid conjugation to the AG. Targeted experiments indicate a specific role for MSMEG0834 in mediating the interaction between MmpL3 and its essential co-factor TtfA. The applicability of our findings was not limited to Mycobacteria: the phenotypic correlation and/or predicted binding between the widely conserved UPF0182-family protein *MSMEG1959, ftsH*, and FtsH regulators suggested a role for this previously enigmatic protein family in membrane protein homeostasis and/or the regulation of FtsH activity. A role in membrane protein homeostasis for UPF0182-family proteins reconciles the diverse phenotypes observed for knockouts in *M. smegmatis*, *M. tuberculosis, C. diphtheriae*, and *B. breve*. Together, these discoveries highlight the number of undiscovered and important gene-gene and gene-pathway connections that can be revealed by broad chemical screens in *Mycobacteria*.

Second, our studies revealed previously under-appreciated effects of first- and second-line antibiotics against *M. tuberculosis* and discovered a novel scaffold-target pair. Ethambutol is a first-line drug for the treatment of tuberculosis. It works by inhibiting EmbABC, enzymes that extend the arabinose chains of AG and LAM using the DPA. Our screen found that, in addition to *embAB,* knockdowns of the sole essential DedA protein were also sensitized. DedA recycles the decaprenyl-phosphate (DP) lipid carrier shared by AG, PG, and PIM synthesis, suggesting that ethambutol causes additional cell stress by sequestering this essential lipid carrier as DPA. We supported this hypothesis by validating a predicted synergy between ethambutol and amphomycin, a drug that further restricts DP availability. D-cycloserine is a second-line drug that primarily targets Alr and Ddl, two enzymes involved in the synthesis of the D-Ala-D-Ala moiety of the PG peptide stem. Our data suggests that in bacteria that express D-alanine transaminase and therefore couple the D-Ala and D-Glu pools, Alr inhibition by D-cycloserine also affects the level of D-Glu, another essential PG constituent. Finally, our screen revealed that PanB is the likely target of salicylic acid, a known antimicrobial without an established mechanism. While aspirin (SA) is unlikely to be an effective treatment for tuberculosis, the discovery of this scaffold-target pair opens the door to further optimization. These findings underscore the complexity of drug mechanisms and point towards novel potential synergies and strategies.

Finally, the phenotypes and correlations in our data will serve as the jumping-off point for numerous future studies in both Mycobacteria and other Actinobacteria. Our screen identified specific phenotypes and correlations for numerous genes that were not further tested. For example, knockdown of *MSMEG0370* (a MukB-like SMC protein associated with efficient plasmid transformation^79^) causes sensitivity to numerous detergents and knockdown of ergothionine synthesis causes sensitivity to cholate, a component of bile (Table S2). Further study of these phenotypes can reveal links between DNA segregation and the cell envelope, and may uncover the physiological role of ergothioneine in diverse gut microbes^80^. In addition to specific phenotypes, our data provide global insights into mycomembrane organization. In gram-negative bacteria, knockdown of lipopolysaccharide synthesis or transport compromises the barrier function of the outer membrane, leading to numerous chemical sensitivities^10^. In contrast, knockdown of mycolic acid synthesis results in both resistance (e.g., to detergents) and sensitivity (e.g., to vancomycin, Table S2). The same is true when targeting other mycomembrane lipids, supporting a previously proposed^81^ model for mycomembrane function in which the tightly packed and covalently attached mycolic acids of the inner leaflet form the primary barrier for lipophilic compounds and outer leaflet lipids, whose properties vary, absorb different lipophilic compounds.

Our work provides a robust foundation for future broad chemical screens, both in *M. smegmatis* and *M. tuberculosis.* Beyond the set of membrane-focused chemicals tested here, we envision performing additional screens targeting other core processes, such as DNA replication, translation, and energy metabolism, to identify novel Mycobacterial and Actinobacterial genes in these essential processes. Given the rich information in our present screen and numerous precedents for the utility of chemical genetic approaches in other bacteria, such efforts will have a transformative effect on the study of Mycobacteria.

## ACKNOWLEDGEMENTS

We thank Jeff Cox and members of the Gross and Rock Labs for extensive helpful discussions. We thank Google DeepMind and Isomorphic Labs for providing free online access to AlphaFold3. We thank Stephanie Bueler and John Rubinstein for their assistance and advice for performing ORBIT and for sharing materials. Work in the Gross lab was supported by National Institutes of Health (NIH) grant R35 GM118061 (CAG). Sequencing was performed at the UCSF CAT, supported by UCSF PBBR, RRP IMIA, and NIH 1S10OD028511-01 grants. NH was supported by a UCSF Pulmonary Division Training Grant from NIH (2T32HL007185), the Burroughs Wellcome Fund Postdoctoral Enrichment Program (1019894), and the University of California President’s Postdoctoral Fellowship Program. NYS was supported by a Medical Scientist Training Program grant from the NIH-NIGMS (T32GM152349) to the Weill Cornell/Rockefeller/Sloan Kettering Tri-Institutional MD-PhD Program. BB was supported by RUCCTS Grant #UL1 TR001866. Work in the Chen Lab was supported by the NIH-NIAID 1DP2AI184740-01 (JC). Work in the Kiessling Lab was supported by the NIH-NIAID AI126592 (LLK) and NSF-GRFP 2141064 (TCW). Work in the Mancia Lab was supported by NIH-NIGMS grant R35 GM132120 (FM). Work in the Glickman Lab was supported by NIH grants P30 AI168433 (Tri-I TRAC).

## METHODS

### Mycobacterial cultures

All *M. smegmatis* strains used in this study were derivatives of mc^2^155.

Unless otherwise specified, liquid cultures of *M. smegmatis* were grown using 7H9 culture medium (BD #271310) supplemented with 0.025% Tween-80 detergent at 37 °C, cultured at 37°C in a rotating drum or shaking at 130RPM. *M. smegmatis* colonies were cultured on 7H9 agar plates at 37 °C. Where required, antibiotics or inducers were used at the following concentrations: kanamycin (KAN) at 20 μg/mL; anhydrotetracycline (ATc) at 100 ng/mL.

### Expansion of the CRISPRi library

Frozen 1mL aliquots of the RLC0011 library (ref) were expanded as follows: 1 mL frozen aliquots were diluted in 7H9 medium + 0.05% Tween-80 + 20 µg/mL KAN at 37 °C. The library was allowed to expand overnight (∼18 hours) to OD600 ∼1.0, then was passed through a 10 µm strainer before being stored at −80 °C in 1mL aliquots.

### Pooled CRISPRi screen

To grow the preculture of the library, a frozen 1mL aliquot of expanded RLC001 *M. smegmatis* CRISPRi library (ref) was added to either 19 mL or 49 mL of 7H9 medium + 0.025% tween + 20 µg/mL KAN and grown at 37 °C overnight to OD600 ≤ 0.8. At this point, a t0 sample was taken.

To initiate the screen, the preculture was diluted to OD600 0.05 in 3mL of 7H9 medium with standard Tween-80 and kanamycin supplements (see *Mycobacterial cultures*), plus 100 ng/μL ATc to induce the CRISPRi system and the stress condition. Pooled CRISPRi screens were performed in deep 24-well plates. Samples were grown for 18 hours (approximately 10 doubling times of WT *M. smegmatis* without stress) before harvesting via centrifugation at 1500 RPM. Samples were resuspended in 75 μL of 1x PBS and stored at −80 °C until DNA extraction.

### Genomic DNA extraction and library preparation for Illumina sequencing

To extract genomic DNA from samples, samples were first lysed via mechanical disruption. Briefly, samples were transferred to a deep 96 well plate with 180 μL of enzymatic lysis buffer (20mM Tris-Cl pH 8.0, 2mM EDTA, 1.2% Triton X-100, 20mg/mL lysozyme) and 0.1 mm silica beads per sample. Samples were subjected to 3 x 1-minute disruption cycles (Qiagen TissueLyser), with 5 minutes of incubation on ice between cycles.

Following lysis, samples were transferred to a fresh 96 well plate, and genomic DNA was extracted via the Qiagen DNeasy 96 Blood and Tissue kit with the Gram Positive Bacteria pretreatment protocol. Genomic DNA concentration was quantified via Qubit BR fluorescence (Thermo Fisher HS kit) measured by plate reader (Biotek).

Amplification of the sgRNA-encoding region was performed via two-step PCR on 25 ng of gDNA. The first step used staggered primers to introduce base diversity during Illumina sequencing and added the TruSeq Adaptors (Table S5). The second step used TruSeq DNA Unique Dual Index Adapters (Illumina), which added i5 and i7 barcodes to each sample and the P5 and P7 flow cell attachment sequences. The first PCR cycling conditions were: 98°C for 45 s; 17 cycles of 98 °C for 10 s, 65 °C for 20 s; 65°C for 300 s. The second PCR cycling conditions were: 98°C for 30 s; 9 cycles of 98 °C for 10 s, 65 °C for 75 s; 65°C for 300 s. The products of PCR1 were cleaned using ExoCIP (NEBiolabs), and 6 μL of cleaned products were used directly as the template for PCR2.

Amplicons were purified via Sera-Mag beads (Cytiva) with one-sided selection (1.8 x). Bead-purified amplicons were visualized and quantified using an Agilent 2100 Bioanalyzer (high-sensitivity chip; Agilent Technologies #5067-4626). Amplicons were multiplexed into 10 nM pools and sequenced on an Illumina sequencer according to the manufacturer’s instructions (5% PhiX spike-in; PhiX Sequencing Control v3; Illumina # FC-110-3001). Samples were run on the NovaSeq 6000 platform with an S4 flow cell (Paired-end 2×150 cycles). Samples were sequenced to achieve a target sgRNA median count depth of approximately 100-150 per sample.

### Sequencing and quantification

Raw FASTQ files were aligned to the library oligos and counted. Reads were summed across replicates of the same condition and relative fitness (RF) was calculated for each sgRNA with at least 100 reads at t0, as previously described^13^. RF was normalized for differences in generation times across experiments as previously described^85^. To facilitate downstream analysis, the RF of sgRNAs targeting each gene were averaged thus: for genes targeted by 10 or fewer sgRNAs, the RF of all sgRNAs was averaged; for genes targeted by more than 10 sgRNAs, sgRNAs are binned into low (0-0.33), medium (0.33-0.66), and high (0.66-1) knockdown strength bins based on their predicted knockdown strength from ref^4^. S-scores are calculated for each gene-bin as previously described^14^.

### Generation of correlation network

For a given pair of genes, the maximum correlation coefficient between the S-scores of their three knockdown bins was calculated. Edges were mapped between two genes if the maximum correlation coefficient between two genes > 0.7. Mapping was done using Cytoscape and the R package RCy3^86^.

### Generation of individual CRISPRi strains

Individual CRISPRi strains were generated as detailed in (Bosch ref^4^). Briefly, the CRISPRi plasmid backbone was digested with BsmBI-v2 (NEB #R0739L). For each individual sgRNA, two complementary oligonucleotides with appropriate overhangs were annealed and ligated (T4 ligase NEB # M0202M) into the BsmBI-v2 digested plasmid backbone. A list of sgRNA sequences used for constructing individual CRISPRi strains can be found in Table S4.

Individual CRISPRi plasmids were electroporated into mycobacteria, recovered, and plated. For each transformation, 100 ng plasmid DNA and 200 μL of electrocompetent mycobacteria were used. For sgRNAs targeting MSMEG3144 and MSMEG3145 (*ripA*), plasmids were co-transformed with pIRL19 (L5 integrase suicide vector^4^).

### Generation of MSMEG3144 complementation plasmids and strains

*MSMEG3144* CRISPRi complementation constructs were cloned as described previously^87^ into the Tweety-integrating backbone plRL91^88^. Each construct was assembled from PCR-amplified fragments using NEBuilder HiFi DNA Assembly Master Mix (NEB #E2621). Fragments were amplified (Q5 High-Fidelity 2X Master Mix; NEB #M0492) from *M. smegmatis* genomic DNA using custom oligonucleotides designed to introduce 20 bp assembly overlaps and, for CRISPRi-resistant constructs, silent mutations in the *MSMEG3144* sgRNA-binding sequence and adjacent PAM site (Table S4). Gel-purified fragments were incorporated into DraIII-digested and gel-purified plRL91 (0.02 pmol backbone, 2–5X fragment excess, 60-minute incubation). To preserve endogenous regulation, all constructs included the intergenic sequence upstream of *MSMEG3144*; the *MSMEG3144* + *MSMEG3145* construct additionally contained the sequence between these two genes. Construct sequences were verified by Sanger sequencing. Validated plasmids were co-transformed with plRL62 (Tweety integrase suicide vector^88^) into *M. smegmatis*, and transformants were selected with nourseothricin (25 μg/mL).

### Time-lapse microscopy

CRISPRi strains were grown to mid-logarithmic phase (OD600 ≈ 0.2) in the absence of inducer, diluted to OD600 = 0.01, and loaded into microfluidic plates (MilliporeSigma #B04A-03) for time-lapse imaging. Upon loading, cells were exposed to a continuous flow of 7H9 medium supplemented with ATc (100 ng/mL) using the CellASIC ONIX2 system (MilliporeSigma #CAX2-S0000) and maintained at 37 °C in an environmental chamber. Phase-contrast images were acquired every 15 minutes for 24 hours using an inverted Nikon ECLIPSE Ti2 microscope with a 60X objective and processed using NIS-Elements AR (Nikon) and Fiji.

### Spot assays

Complementation strains were synchronized in logarithmic phase (OD600 ≈ 1), diluted to OD600 = 0.17 (∼50,000 cells/μL), and subjected to four successive 10-fold dilutions to generate comparable five-point dilution series. Each series was plated using a multichannel pipette (2 μL per dilution) onto 7H10 agar containing nourseothricin (25 μg/mL), with or without ATc (100 ng/mL). Plates were incubated at 37 °C and imaged after approximately 50 hours (∼20 generation times).

### AzLac labeling assays

#### Flow cytometry

*M. smegmatis* was cultured in Middlebrook 7H9 containing 0.05 % Tween 80 to OD600 0.2. Once this density was reached, the CRISPRi system was induced with 100 ng/mL ATc for 5 hours, at which point the sample was diluted to 0.1x concentration in 7H9 + 0.05% Tween 80, and 100 μL of this suspension was transferred to a 96-well plate. For treatment samples, 100 μM AzLac^26^ was added. Unless otherwise stated, the final concentration of AzLac was 100 µM. Cultures were grown for 16 hours at 37 °C shaking. Samples were immediately prepared for flow cytometry.

The preparation of flow cytometry samples was performed as previously described^26^. Briefly, cells were pelleted for 5 min at 3000 xg, washed once with ice-cold phosphate-buffered saline (PBS) supplemented with 0.05% Tween 80, then washed an additional time with PBS supplemented with 0.05% Tween 80 and 0.5% (w/v) bovine serum albumin (BSA) before being resuspended in fresh 7H9 media supplemented with 0.5% Tween 80. AFDye™ 647 DBCO (Click Chemistry Tools #1302) was added to a final concentration of 250 µM. The samples were stained for 2 hours rotating at 37 °C.

Stained cells were pelleted for 5 min at 3000 x g. The supernatant was removed, and the pellet was washed twice with PBS supplemented with 0.05% Tween 80 and 0.5% (w/v) BSA. Stained cell pellets were fixed in 4% paraformaldehyde in PBS for 20 min. Following fixation, cells were resuspended in PBS supplemented with 0.05% Tween 80 in flow tubes and analyzed using an Attune NxT Flow Cytometer. 10,000 cells were counted at the low flow rate. Data were analyzed using the FlowJo software package (FlowJo LLC). Mean fluorescence intensity was calculated using a geometric mean.

### Quantification of arabinan/galactan ratio

#### mAGP Isolations

The mAGP was isolated according to ref^89^. Briefly, 1 mL of unlabeled *M. smegmatis* was aliquoted at a normalized optical density (OD600 = 1.0-2.5). Aliquots were centrifuged for 5 minutes at 3000 xg at 4 °C and the supernatant was removed. 1 mL of 2% Triton-X100 in PBS was added to the pellets, and the resulting suspensions were sonicated for 2 minutes and repeated for a total of 5 cycles. The suspensions were then centrifuged at 15,000 xg for 15 minutes. The supernatant was removed and the pellet was resuspended in 1 mL of 2% SDS in PBS. The suspension was then heated to 95 °C for 1 hour, followed by centrifugation at 15,000 xg for 15 minutes. Using the same centrifugation steps, the pellet was washed with water, 20% water in acetone, and 100% acetone. The pellet was then dried overnight.

### Hydrolysis and Composition Analysis of the mAGP

After isolation, mAGP pellets were resuspended in 150 µL of 18% trifluoroacetic acid in water (v/v) and left at room temperature. After letting the hydrolysis reaction proceed, the solution was diluted with 350 µL water and was filtered through a 0.22 µM syringe filter. The solvent was then removed via rotary evaporation under reduced pressure. To this solid, 150 µL of a solution of 0.25 M sodium borohydride and 1 M ammonia in ethanol was added and left to react overnight on a rocker. After 18 hours, 150 µL 10% acetic anhydride in methanol (v/v) was added and left to react for 2 hours. The solutions were concentrated under reduced pressure. 500 µL of 50% acetic anhydride in pyridine (v/v) was added to the solid, and the mixture was placed on a rocker overnight. After 18 hours, 500 µL of toluene was added and the solutions were concentrated under reduced pressure. 1 mL of ethyl acetate was added to the solid and the resulting solution was filtered through a 0.22 µM syringe filter. The solution was concentrated under reduced pressure, the samples were dried under a vacuum, and the residue was placed in a −20 °C freezer for storage. The samples were redissolved in 1:1 LCMS grade acetonitrile:water. Mass spectrometry was run on an Ultivo triple-quadrupole coupled to an Agilent 1260 Infinity II LC system eluting with a gradient from 17 to 23% acetonitrile in water over 25 min with a flow rate of 0.1 mL/min.

### Generation of correlation network

For a given pair of genes, the maximum correlation coefficient between the S-scores of their three knockdown bins was calculated. Edges were mapped between two genes if the maximum correlation coefficient between two genes > 0.7. Mapping was done using Cytoscape and the R package RCy3^86^.

### Generation of endogenously tagged mycobacteria strains

Endogenously tagged *M. smegmatis* strains were generated using ORBIT^36^. All strains were grown in Middlebrook 7H9 media (Difco) (supplemented with 0.5% w/v BSA, 11 mM dextrose, 14 mM NaCl, 0.2 % w/v glycerol, 0.05% w/v Tween-80), and appropriate antibiotics. An initial strain was electroporated with pKM461 (Addgene #108320), which allows for the expression of a Che9c phage annealase and a Bxb1 integrase. pKM461 was a gift from Kenan Murphy. A single colony of the pKM461 containing strain was used to inoculate a 10 mL starter culture which was grown to saturation at 37 °C in a shaking incubator at 180 RPM. This starter culture was used to inoculate a second 25 mL culture which, at an OD600 0.5, was supplemented with 500 ng/mL anhydrotetracycline (SupelCo #37919) to allow protein expression. The culture was grown at 37 °C for another 3.5 hours. 20 mL of this culture was harvested, and the cells were pelleted by centrifugation at 3500 RPM. The pellet was then washed twice in sterile, ice cold 10% glycerol. For strain generation, 380 μL of electrocompetent cells were mixed with 200 ng of the payload plasmid pSAB41, a derivative of pKM491 (Addgene #109282) containing a 3xFLAG tag, and 1 μg of a targeting DNA oligomer (Table S4). pSAB41 was a gift from Stephanie Bueler and John Rubinstein. The cells were then electroporated in an ice-cold 0.2 cm gap cuvette (BioRad #1652086) and allowed to recover in 2 mL antibiotic free Middlebrook 7H9 media overnight. 200 μL of the culture was then plated on a 7H10 plate containing 50 μg/mL hygromycin B (Goldbio #H-270-1) and incubated at 37 °C for 4 to 5 days. Positive colonies were verified for the chromosomal insertion using PCR or whole genome sequencing (Plasmidsaurus).

### Expression and purification of chromosomally tagged proteins

Strains were grown in Middlebrook 7H9 media (Difco) supplemented with 11 mM dextrose, 14 mM NaCl, 0.2% w/v glycerol, 0.05% w/v Tween-80 and 50 μg/mL hygromycin B. The 3x-FLAG tagged AftC and the 3x-FLAG tagged MSMEG3144 strain were each streaked on LB agar plates containing 50 μg/mL hygromycin B and incubated at 37 °C for 2 days. Colonies were picked and used to inoculate a 20 mL starter culture which was grown at 37 °C at 180 RPM for two days. The following day, this culture was used to inoculate 3 L of Middlebrook 7H9 media and grown at 37 °C shaking at 180 RPM for two days. The cells were then harvested with centrifugation at 6000 x *g* for 10 minutes in an Avanti J-E centrifuge (Beckman Coulter) and flash frozen until further use. After thawing, the cells were resuspended in Lysis Buffer (50 mM Tris pH 7.5, 100 mM NaCl, 20 mM MgSO_4,_ and 5 mM TCEP) using 5 mL buffer per gram of cells. The solution was then supplemented with 20 μg/mL DNase1, 0.2 μg/mL phenylmethylsulfonyl fluoride (PMSF), and EDTA-free complete protease inhibitor cocktail (1 tablet/liter) (Roche) and lysed with 4 to 5 passages through an Emulsiflex C3 homogenizer (Avastin) operating at a pressure of 10,000 – 15,000 PSI. Cell debris and unlysed cells were pelleted by centrifugation at 3000 x *g* in a Centrifuge 5510 R (Eppendorf) for 10 minutes.

Membranes were then pelleted by centrifugation at 185,600 x *g* in a Type 45 Ti Rotor (Beckman Coulter) for 1 hour at 4 °C. The membrane pellets were resuspended in Lysis Buffer (15 mL/gram of membrane) and flash frozen until further use. After thawing, n-docyl-β-D-maltopyranoside (DDM) (Inalco #1758-1350) was added to the solution to a final concentration of 1% and stirred at 4 °C for 3 hours. The solution was then clarified with centrifugation at 185,600 x *g* for 30 minutes at 4 °C. The supernatant was then applied to a gravity column packed with 200 μL of α-FLAG G1 Affinity resin (GenScript #L00432). The column was washed with 10 CV of Wash Buffer 1 (50 mM Tris pH 7.5, 150 mM NaCl, and 0.02% DDM) and 10 CV of Wash Buffer 2 (50 mM Tris pH 7.5, 300 mM NaCl, and 0.02% DDM). Protein was eluted with 4 CV of Wash Buffer 1 containing 150 μg/ mL of 3xFLAG peptide (GLPBIO #GP10149). The elution fraction was then concentrated 4-fold using an Amicon Ultra Centrifugal Filter (Millipore Sigma #UFC501024) and 10 μg of protein was mixed with 6x reducing Laemmli SDS Sample Buffer (Boston BioProducts #BP-111R) and analyzed with SDS-PAGE using 4-20% Mini-PROTEAN TGX Precast Protein Gels (BioRad #456109), followed by staining with Instantblue Coomassie Protein Stain (Adcam #ab119211).

### Mass spectrometry protein identification

After SDS-PAGE analysis, bands of interest were excised and transferred to 20 μL of sterile 1% acetic acid and stored on dry ice. Mass spectrometry analysis was performed by The Protein Facility of the Iowa State University Office of Biotechnology. The samples were thawed and reduced with DTT, the cysteine residues were modified with iodoacetamide, and digested overnight with trypsin/Lys-C. The digestion was stopped with the addition of formic acid, and the samples were dried in a SpeedVac. Samples were then desalted using a BioPureSPN MiniPROTO 300 C18 column (Nest Group Bio) before drying again in a SpeedVac. PRTC standard (Pierce #88320) was spiked into the sample to serve as an internal control. The peptides were then separated by liquid chromatography on an EASY-Spray PepMap Neo 75 μm/150 μm x 150 mm column (ThermoScientific #ES75150PN), operated on a Vanquish NEO HPLC system coupled with an Easy-Spray Source (ThermoScientific). The column was washed with Buffer A (0.1% formic acid) and peptides were eluted with Buffer B (80% acetonitrile, 0.1% formic acid). The peptides were analyzed by MS/MS on an Orbitrap Astral (ThermoScientific) by fragmenting each peptide. The resulting intact and fragmentation pattern was compared to a theoretical fragmentation pattern to find usable peptides for protein identification. The PRTC areas were used to normalized data between samples. The resulting peptides were searched using CHIMERYS^90^ against the *M. smegmatis* mc^2^155 genome and analyzed with ProteomeDiscover (ThermoScientific).

### MSMEG0834 knockout in merodiploid

To create truncation strains of MSMEG0834, a deletion of *MSMEG0834* was made by homologous recombination and double negative selection^91^ in a mc^2^155 strain containing a second copy of *MSMEG0834* at *attB(L5)*. The second copy could encode a truncated MSMEG0834.

### TtfA-msfGFP pulldown with CRISPRi depletion

TtfA-msfGFP pulldowns were done as previously described^32^ with the following modifications. 40mL cultures of TtfA-msfGFP strains with CRISPRi targeting constructs were grown to an OD600 of 0.5 in LB supplemented with 0.5% glycerol + 0.5% dextrose overnight. 25µL of Lag16-2-agarose (see “Lag16-2 purification and agarose coupling”) was used per sample for GFP pulldown. Elution was performed using SDS sample buffer with heating at 50°C for 15 min.

### Lag16-2 purification and agarose coupling

Lag16-2 expressed and purified as described^92^ with the following modifications. A 19 hour induction was done at 22°C in BL21(DE3) cultured in terrific broth containing carbenicillin (100 μg/mL). Eluates were pooled, concentrated in Amicon Ultra-4 3k, and run on Superdex 200 10/300 size exclusion column using an AKTA purifier in phosphate-buffered saline (PBS). 8 mg of purified Lag16-2 in 5ml of PBS was added to 300mg of NHS-Activated Agarose (Pierce, PI26196) and coupled by mixing 18 hours at 4 °C. Agarose was collected by centrifugation at 500g for 1min at 4 °C then washed with 15ml TBS. Agarose was collected by centrifugation at 500g for 1min at 4 °C, supernatant was discarded and resin was quenched in 15mL TBS for 30min at 4°C. Quenched Lag16-2-agarose was washed 6 times with 15mL PBS with a final suspension of 4ml total in PBS at ∼2mg/mL Lag16-2 in agarose slurry.

### Immunoblotting

For protein and epitope tag detection, the following antibodies were used: GFP (rabbit anti-GFP polyclonal antibody, 1 mg/mL, 1:20,000; Rockland Immunochemicals), mCherry (rabbit anti-RFP polyclonal antibody, 1 mg/mL, 1:20,000; Rockland Immunochemicals), VirR (rabbit sera, 1:10,000)^29^.

### Bacterial adenylate cyclase-based two-hybrid (BACTH)

Plasmids derived from pUT18 and pKT25 containing T18 and T25 fusions to mycobacterial proteins were transformed into a *cya-E. coli* strain, DHMI, [F^−^ *glnV44*(AS) *recA1 endA gyrA96 thi-1 hsdR17 spoT1 rfbD1 cya-854*]^34^ and selected on LB agarose containing kanamycin (40µg/mL) and carbenicillin (100µg/mL). Resulting transformants were patched on LB agarose containing kanamycin, carbenicillin, and X-gal (40µg/mL), and IPTG (0.5mM) and incubated overnight at 37°C.

## DATA AVAILABILITY

Raw sequencing data is deposited on NCBI-SRA Bioproject PRJNA1367630.

## SUPPLEMENTAL INFORMATION TITLES AND LEGENDS

**Figure S1.**
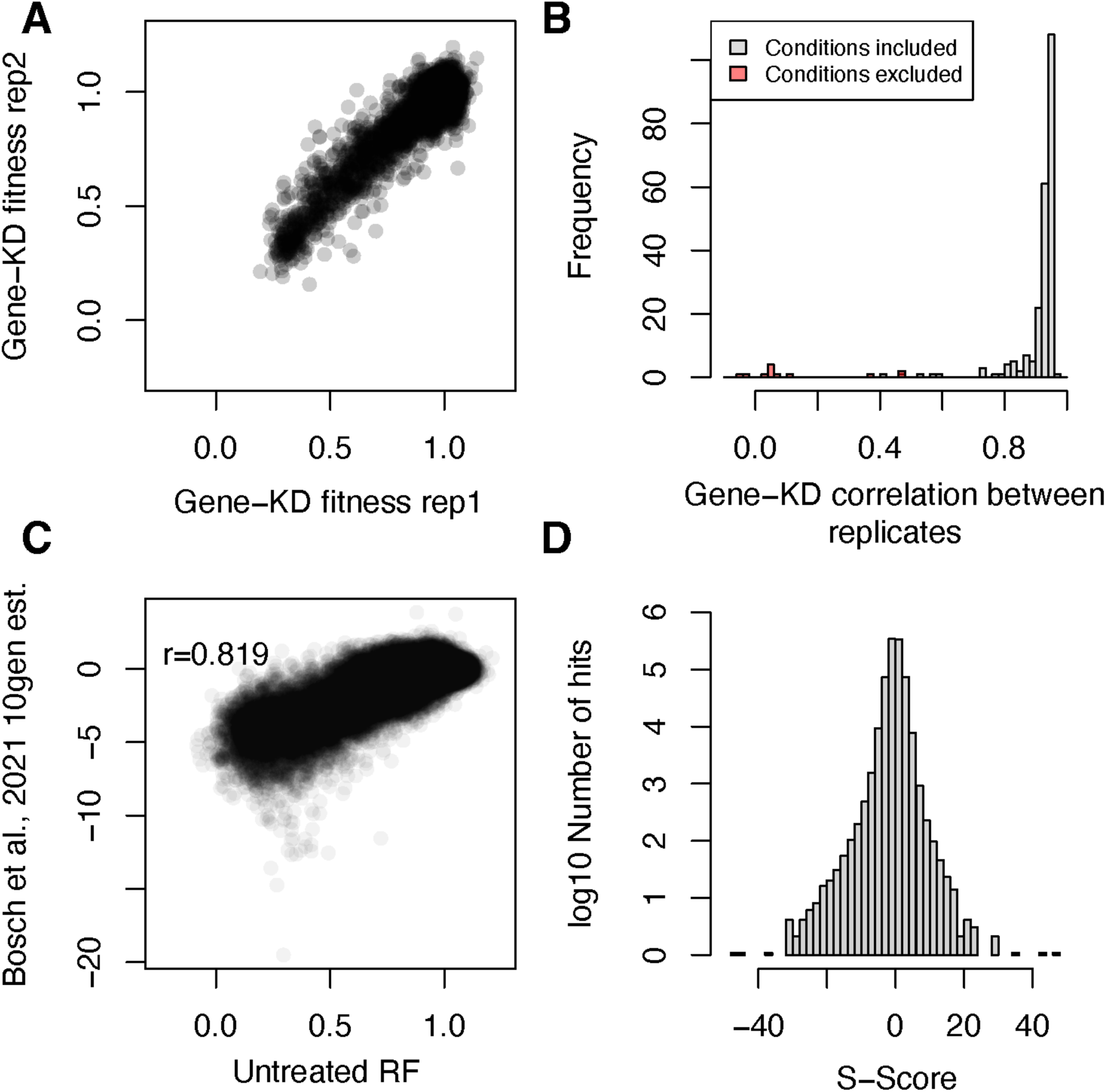
RF was reproducible and consistent with previous measurements of the library. A) Per gene per bin fitness in two replicates of the no-chemical control shows high reproducibility. B) Histogram of per gene per bin reproducibility for all conditions tested. Red bars represent reproducibility of conditions that were excluded. C) per sgRNA fitness of the no-chemical control sample is highly correlated to previously published measurements of the library. D) Histogram of the total number of hits at different S-score thresholds. Note logarithmic y-axis.

**Figure S2.**
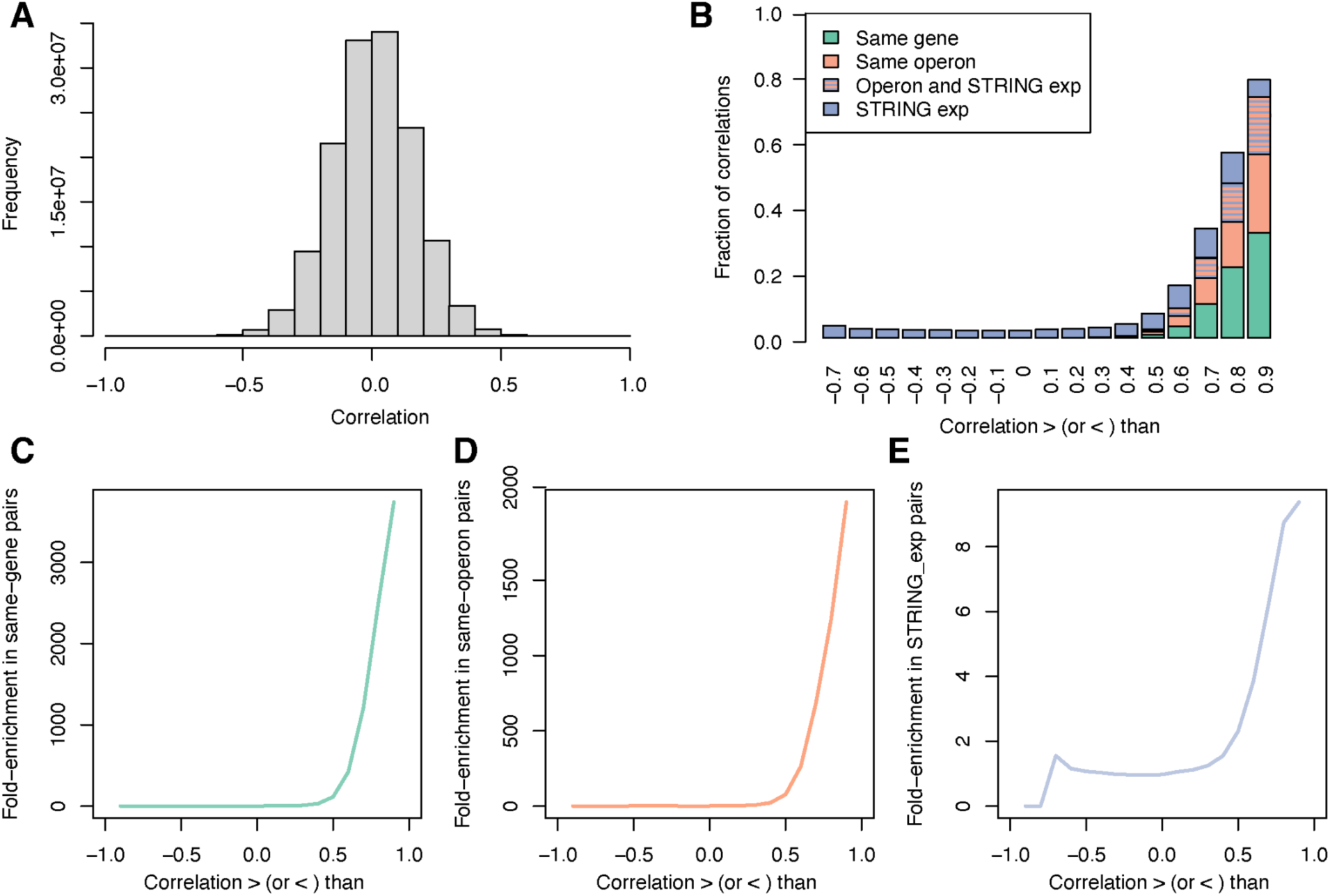
Correlations between the phenotypic profiles of genes are indicative of shared function. A) Histogram of phenotypic correlations between gene-knockdown bins. B) Highly correlated phenotypic profiles include different knockdown levels of the same genes (green), different genes in the same operon (orange and blue), or between separately transcribed genes with some experimental evidence of interaction in the STRING database (purple). Highly correlated phenotypic profiles are enriched in different knockdown levels of the same genes (C), genes in the same operon (D) and genes with experimental evidence of interaction (E).

**Figure S3.**
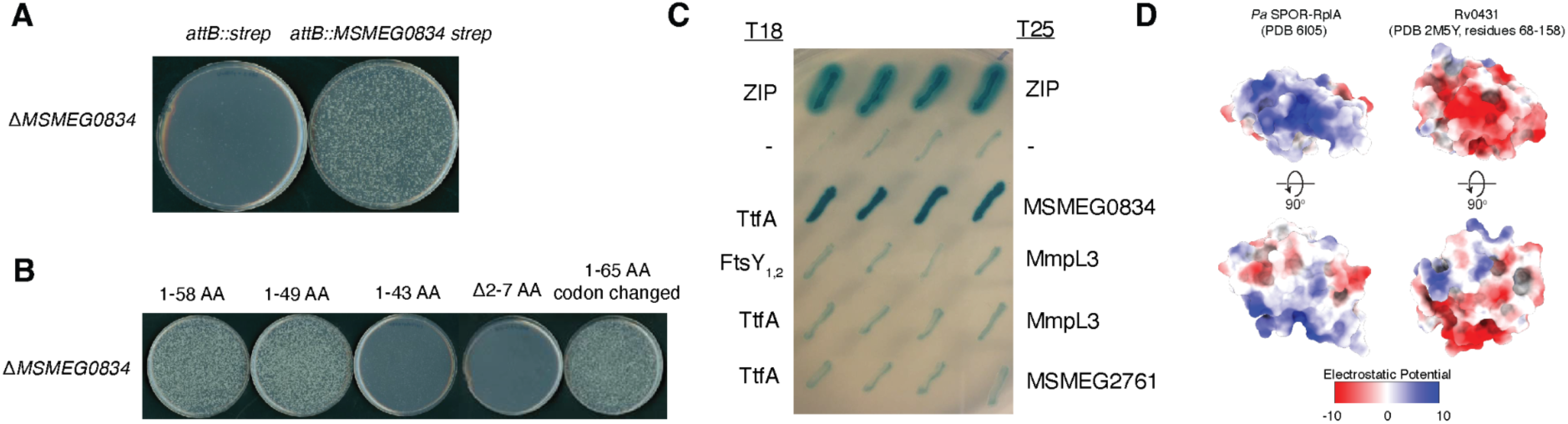
MSMEG0834 is essential and mediates the interaction between TtfA and MmpL3 via its N terminus. A) A Δ*MSMEG0834 attB::MSMEG0834* strain was transformed with allelic exchange *attB*(LB) vectors of either *attB:strep* or *attB:MSMEG0834-strep* to assess essentiality. Transformants without a copy of *MSMEG0834* are unviable. B) A Δ*MSMEG0834 attB::MSMEG0834* strain was transformed with allelic exchange *attB*(LB) vectors of *attB:MSMEG0834[TRUNCATION]-strep* to assess essentiality. Truncation is specified above plates. Truncations as short as the first 49AA of MSMEG0834 are able to support growth. C) BACTH assay between TtfA, MSMEG0834, MmpL3, and other membrane proteins. Of the pairs tested, only TtfA and MSMEG0834 directly interact. D) Electrostatics of the *Pseudomonas aeruginosa* RplA SPOR domain compared to the MSMEG0834 LytR_C domain indicate very different charges, suggesting that the MSMEG0834 LytR_C domain does not bind the same ligand as the SPOR domain.

**Figure S4.**
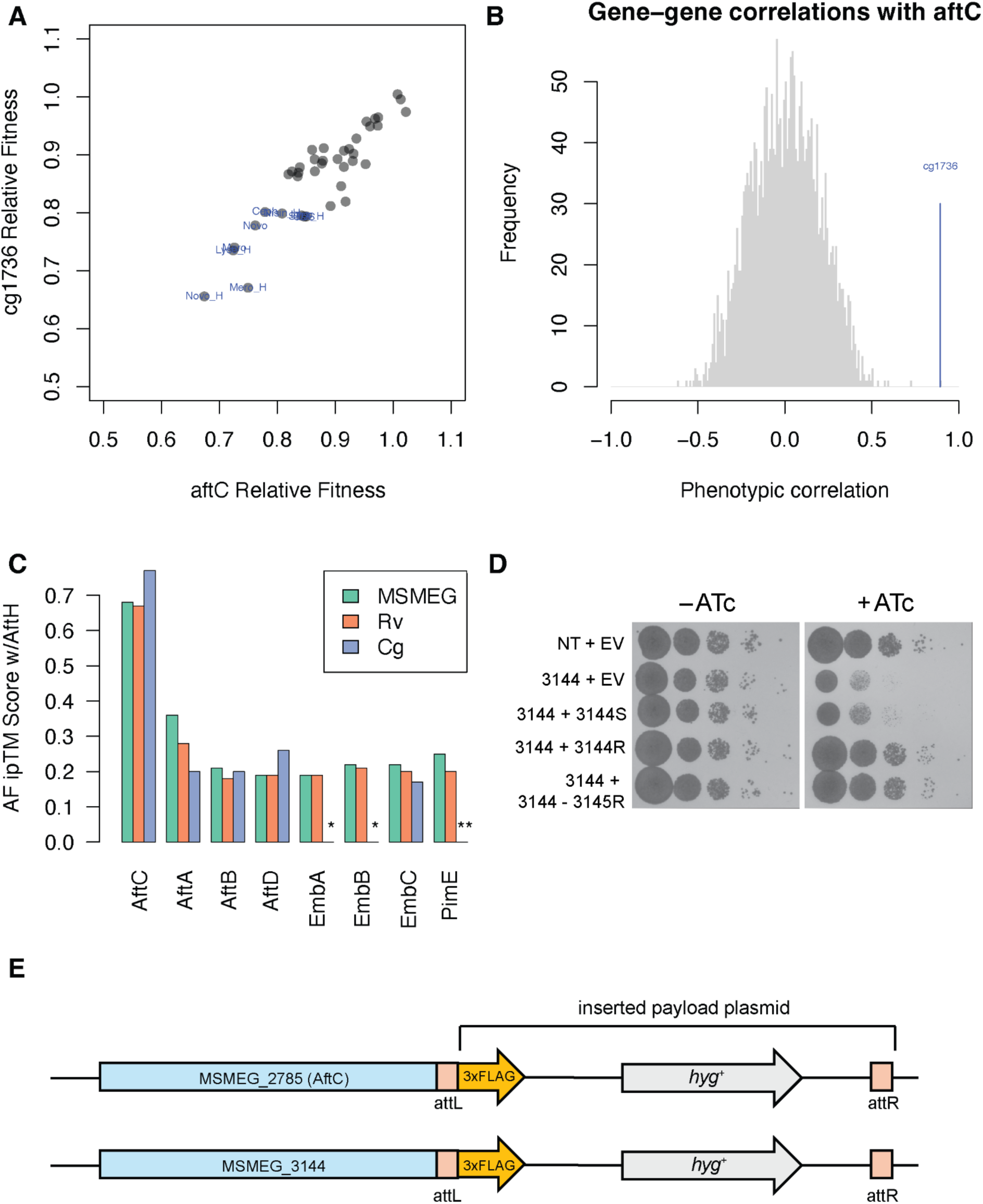
AftC and MSMEG3144 interact to form a functional complex. The phenotypes of *C. glutamicum MSMEG3144* homolog (*cgp_1736*) Tn-insertion mutants is highly (A) and specifically (B) correlated to the phenotypes of the *aftC* mutants in that species^38^. C) AlphaFold3 predicts a high confidence complex between AftC and MSMEG3144 homologs in *M. smegmatis, M. tuberculosis,* and *C. glutamicum.* No significant interaction is predicted between MSMEG3144 homologs and homologs of other arabinosyltransferases. \**C. glutamicum* encodes a single *emb* gene that is most similar to *embC*^83^. \*\**C. glutamicum* does not encode a PimE homolog^84^. D) A CRISPRi-resistant complementation construct carrying *MSMEG3144* alone was sufficient to rescue the growth defect caused by *MSMEG3144* knockdown, indicating that the phenotypes of the *MSMEG3144* knockdown strain are not due to polar effects. NT = non-targeting; EV = empty complementation vector; S = CRISPRi-sensitive (unaltered coding sequence); R = CRISPRi-resistant (silent mutations in PAM and seed sequence) E) Schematic of ORBIT constructs used to create MSMEG3144-3xFLAG and AftC-3xFLAG strains.

**Figure S5.**
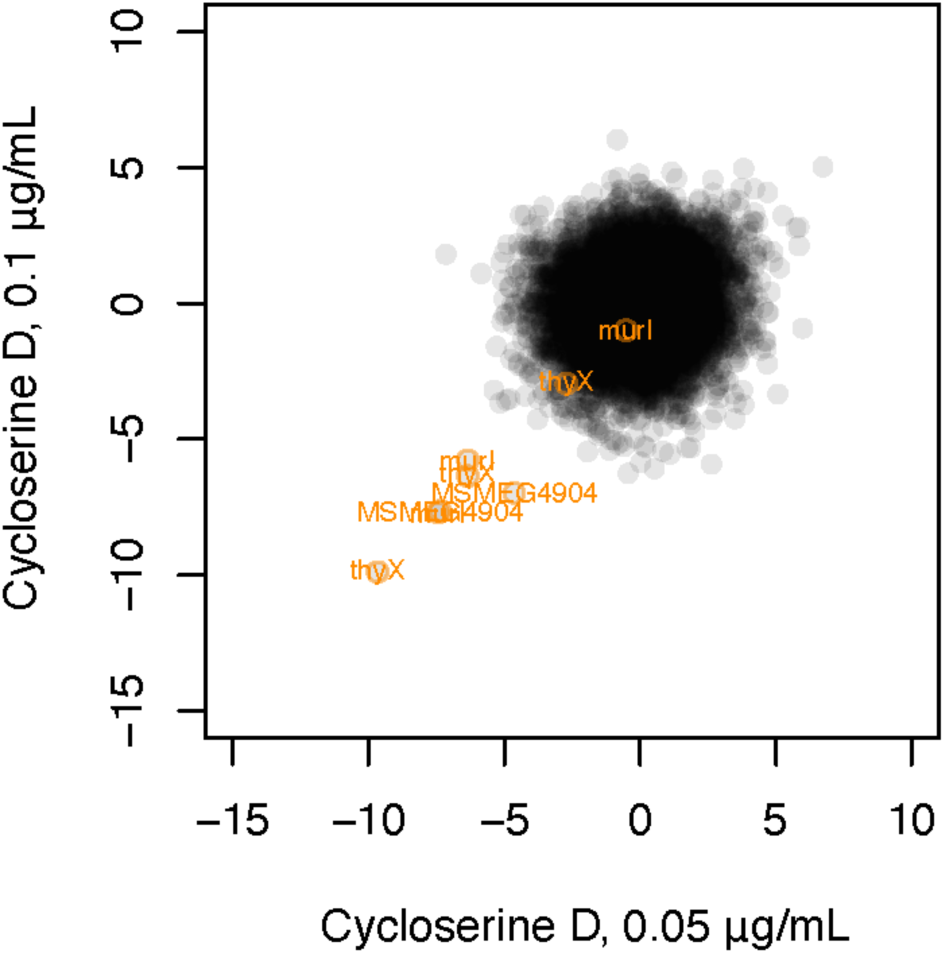
S-scores for all genes and knockdown levels for two concentrations of D-cycloserine. Knockdown strains targeting *thyX*, *murI*, and *MSMEG4904* are sensitized (highlighted in orange).

**Figure S6.**
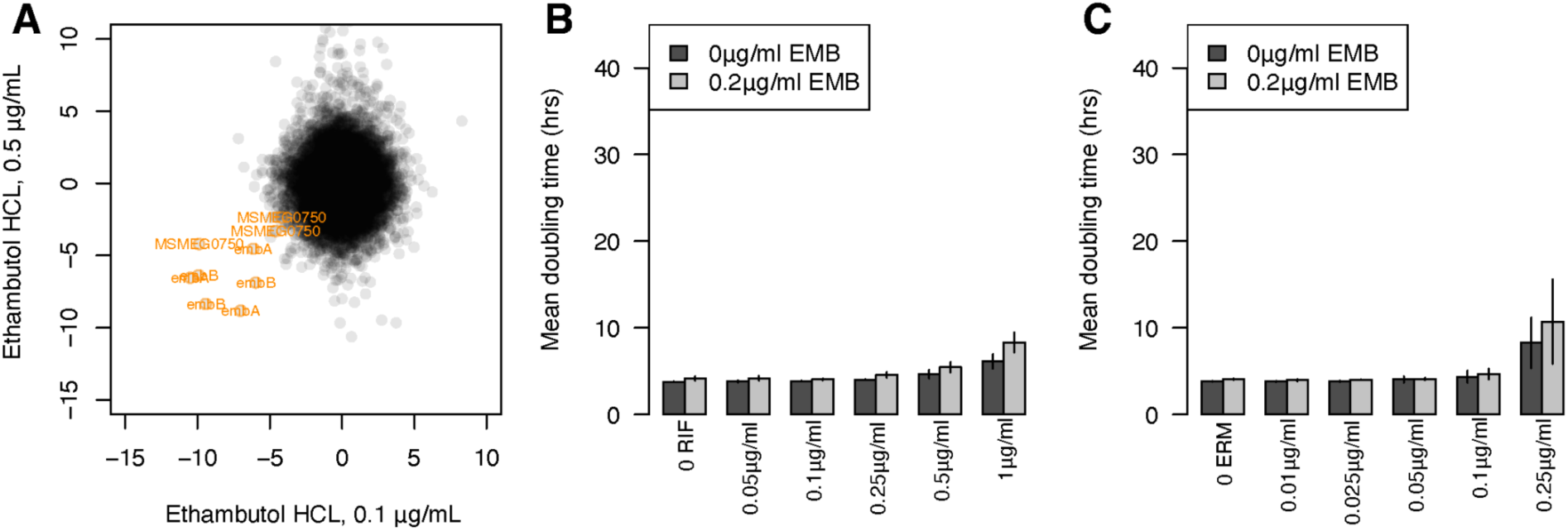
CRISPRi strains targeting *dedA* (*MSMEG0750*) are sensitized to ethambutol. A) S-scores for all genes and knockdown levels for two concentrations of ethambutol. Knockdown strains targeting the essential targets of ethambutol (*embA, embB)* and the essential DedA homolog *MSMEG0750* are sensitized (highlighted in orange). Ethambutol also inhibits EmbC, but EmbC is not essential in *M. smegmatis* (although it is essential in *M. tuberculosis*)^4^. A sub-MIC concentration of ethambutol (0.2µg/ml) did not sensitize cells to rifampicin (B) or erythromycin (C), (compare to Figure 6D).

**Figure S7.**
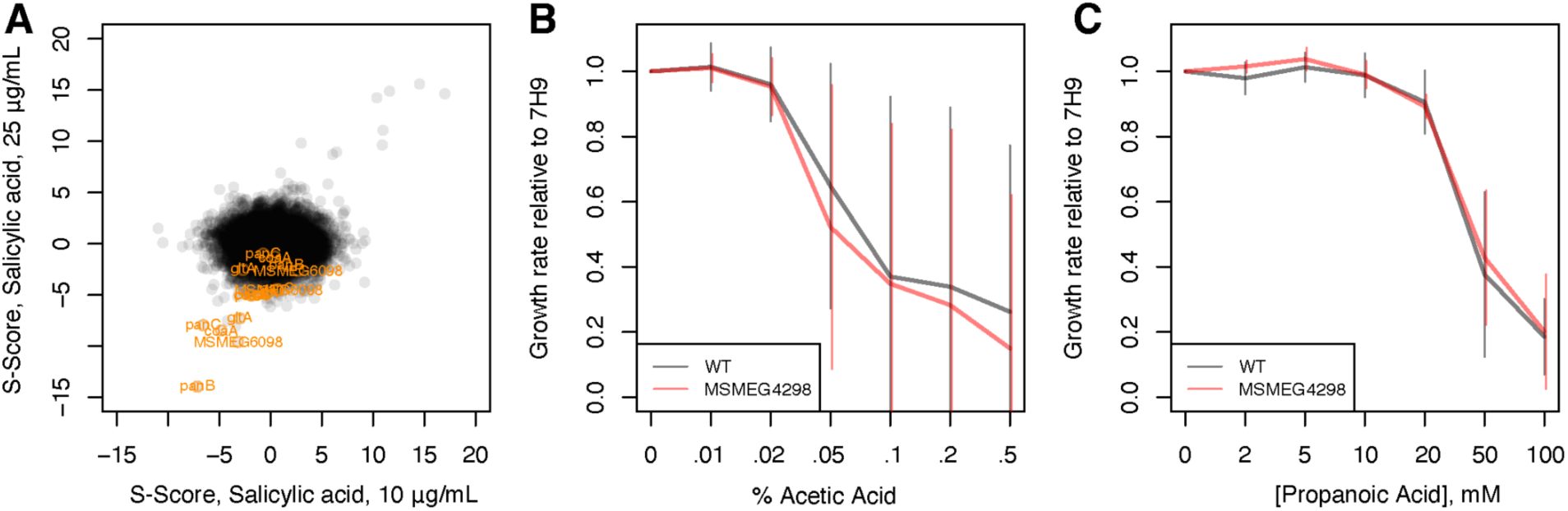
Knockdown strains of early CoA synthesis genes are specifically sensitized to SA. A) S-scores for all genes and knockdown levels for two concentrations of SA. Knockdown strains targeting *panB-MSMEG4298*, *panC-MSMEG6097*, *panG-MSMEG6098*, *coaA-MSMEG5252, sucCD* (succinate-CoA ligase) and *gltA* (citrate synthase) are sensitized (highlighted in orange). The *panB* knockdown strain is not sensitized to either acetic (B) or propanoic acid (C) relative to the WT strain (compare to Figure 7B).

## TABLES WITH TITLES AND LEGENDS

**Table S1** Relative fitness for all strains across all conditions tested.

**Table S2** S-scores for all gene-knockdown levels across all conditions passing quality control.

**Table S3** Correlations between all gene-knockdown levels across conditions.

**Table S4** Primers and strains.

